# Identification of divergent *Toxoplasma* Nuclear Pore Complex components highlights speciation of mRNA export machinery

**DOI:** 10.1101/2025.08.27.672535

**Authors:** Pravin S. Dewangan, Shannon R. Dohr, Jackson T. Trotter, Bailey Nichols, Michael L. Reese

**Affiliations:** Department of Pharmacology, University of Texas, Southwestern Medical Center, Dallas, TX USA; Department of Biochemistry, University of Texas, Southwestern Medical Center, Dallas, TX USA

**Keywords:** Evolutionary cell biology, Nuclear pore complex, Apicomplexa, Parasitology

## Abstract

**Background:** A hallmark of the eukaryotic cell is the regulated transport between the nucleus and cytoplasm, which is mediated by a multi-subunit protein assembly called the nuclear pore complex (NPC). While its overall architecture has been preserved across eukaryotes, the NPC structure varies in different organisms, which appears to have tuned its function. Outside of a handful of model systems, the NPC has not been comprehensively studied. This is particularly true of species that are not closely related to well-studied models, such as apicomplexan parasites. Indeed, the evolutionary divergence of Apicomplexa has complicated facile prediction of NPC proteins in these organisms. Because of this, the NPC components remain largely unidentified, and therefore NPC cellular function in Apicomplexa is poorly understood.

**Principal Findings:** Here we identified, experimentally validated, and functionally characterized protein components of the NPC in the apicomplexan parasite *Toxoplasma gondii*. By combining proximity biotinylation with careful bioinformatic analysis, we identified 16 previously uncharacterized proteins that localize to the *Toxoplasma* NPC. We demonstrated 8 of these proteins are essential to parasite replication. Importantly, we defined components of the mRNA export machinery, as well as Nups required for the stability and/or assembly of specific NPC subcomplexes. Consistent with the evolutionary distance between *Toxoplasma* and well-studied models, the majority of our newly validated NPC components show no clear homology to NPC proteins in yeast, animals, or plants. Moreover, we demonstrated that the *Toxoplasma* mRNA export machinery has a distinct composition from other well-established systems. Intriguingly, Sus1, a well-defined protein of the TREX-2 and SAGA complexes, is missing from the *Toxoplasma* genome. In contrast, others, such as Centrin-3, have been conserved in *Toxoplasma*, but are not required for mRNA export in the parasite.

**Conclusion:** Our work highlights the distinct composition of multiple subcomplexes of the *Toxoplasma* NPC and paves the way for future studies to provide high-resolution structural information on the parasite’s unusual NPC architecture.

## Introduction

The nuclear pore complex (NPC) is one of the largest macromolecular assemblies in a eukaryotic cell and serves as the gateway through which RNA and proteins are trafficked between the nucleus and cytoplasm [1]. In all eukaryotes, the NPC is composed of a series of concentric rings with 8-fold symmetry [2–4]. The NPC is well-conserved across eukaryotic phyla, and structural studies have revealed a common core architecture in plant, fungal, and animal species [5–10]. The NPC rings are composed of multiple biochemically-defined subcomplexes, comprised of proteins called nucleoporins (Nups) [11–14] (Figure 1A), and it is these subcomplexes that are the most highly conserved. Indeed, it has been suggested that the major components of the NPC were conserved in the last common eukaryotic ancestor [15]. Nevertheless, while its core structure is preserved, the NPC varies across different lineages [16,17]. Nowhere is this more evident than in apicomplexan parasites like *Toxoplasma gondii*. A recent cryo-electron tomography study compared the in-cell architectures of the NPC from yeast, human, and *Toxoplasma* cells and found that the parasite’s NPC is unusually asymmetrical, having lost the Y-shaped complex from its cytoplasmic face [7] (Figure 1A). In addition, while the *Toxoplasma* nuclear basket could not be resolved, there appeared to be a much larger membrane-anchored core in the parasite NPC [7] (Figure 1A).

**Figure 1:**
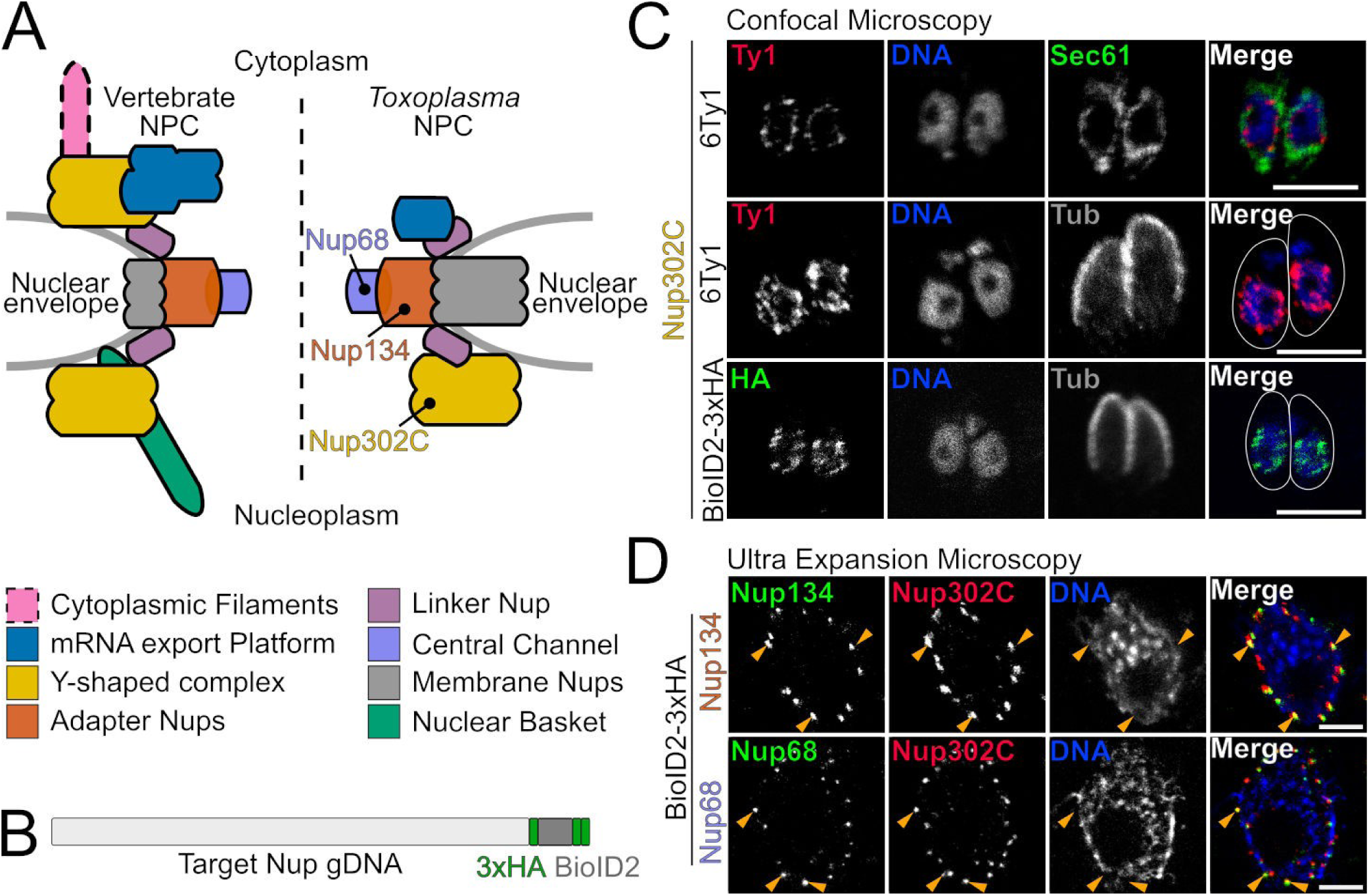
*Toxoplasma* Nup BioID strains localize to NPC. (A) Cartoon overview highlighting differences in the well-studied vertebrate (left) and *Toxoplasma* (right) NPC architectures. The three proteins we use as BioID baits and their respective subcomplexes are indicated by color. (B) Each Nup was endogenously tagged to express an in-frame C-terminal 3xHA-BioID2. (C) Nup302C^6xTy1^ is at the NPC, and present between the cytoplasmic face of the ER (Sec61) and the nucleus (DNA). In all candidate strains, we use Nup302C^6xTy1^ as a marker for the NPC. (D) Both Nup134 and Nup68 localize with Nup302C^6xTy1^ puncta at the NPC by UExM (orange arrows). All scale bars are 5 μm. Note in UExM samples, scale bars represent ∼5× expanded size.

In addition to *Toxoplasma*, the phylum Apicomplexa includes the causative agents of a number of the world’s most devastating diseases, such as the *Plasmodium* species that cause malaria. *Toxoplasma* and other apicomplexan parasites are members of the Alveolate superphylum and are highly divergent from well-studied animal, fungal, and plant models. Because of this, many proteins that are well-conserved in other organisms are difficult to identify by standard bioinformatic methods. This has stymied research into many basic cellular processes, including nucleocytoplasmic transport (NCT) through the NPC. To date, a handful of studies have validated 11 proteins as NPC components in *Plasmodium* [18,19] and 8 in *Toxoplasma* [20,21]. In *Tetrahymena*, *Trypanosoma*, yeast, and humans, the NPC is composed of multiple copies of ∼25-30 proteins [22–25], suggesting that many NPC components in Apicomplexa have yet to be identified and experimentally validated. Components of the NPC are called nucleoporins or Nups and have historically been numerically named based on their apparent molecular weights on an SDS-PAGE (*e.g.* “Nup57”). This means that orthologous proteins rarely share a numerical name and non-orthologous proteins in different organisms can have the same number. To avoid confusion, we will use a numerical Nup nomenclature only when referring to Nups from model organisms, such as yeast and those few parasite proteins that have been previously named.

Here we integrated proximity biotinylation with carefully curated bioinformatic analysis to identify and validate 16 additional *Toxoplasma* Nups. We went on to demonstrate that, as in other organisms, the *Toxoplasma* NPC is robust to deletion of individual Nups, in that only a handful of the proteins we validated as Nups are essential. We also identified components of two NPC-associated subcomplexes that are required for the export of mRNA. Finally, using markers for two distinct NPC subcomplexes that localize to opposing faces of the nuclear envelope, we identified Nups required for either the assembly or stability of NPCs in *Toxoplasma.* Together these data provide the most comprehensive functional characterization of the components of the NPC in an apicomplexan organism and will enable future in-depth cellular and biophysical studies.

## Results

### Coupled bioinformatic analysis and proximity biotinylation identifies components of the Toxoplasma nuclear pore complex

In diverse organisms where the NPC has been most thoroughly investigated, there are approximately 30 Nups that comprise the NPC [22–26]. Previous efforts have predicted components of the *Toxoplasma* NPC by bioinformatic analysis [27] and immunoprecipitation/mass spectrometry [20,21]. To date, 8 of these proteins have been validated as localizing to the NPC [20,21]. Our first goal was to expand this number. Because the NPC is highly stable, we reasoned that proximity biotinylation [28,29] would be an excellent method to identify new Nups in *Toxoplasma*. Because we saw toxicity when fusing some proteins to TurboID [30], we decided to use the smaller BioID2 [31] protein. As probes, we chose 2 validated *Toxoplasma* Nups and 1 additional Nup after validating its localization (Figure 1). These proteins are predicted by their homology to be associated with three different NPC subcomplexes (Figure 1; TgNup68 – Central channel; TgNup134 – Adapter; TgNup302C – Y-shaped complex). Each of these proteins faithfully localized to the *Toxoplasma* NPC when endogenously tagged at their C-termini with a BioID2-3xHA fusion (Figure 1B-D). Note that while TgNup134 and TgNup302 had been previously validated, TgNup68 had not been previously localized to the NPC. We found that both Nup134 and Nup68 showed a [12,32] clear co-localization with Nup302C at the NPC (Figure 1D). For proximity biotinylation experiments, parasites for each BioID strain and the untagged parental strain were grown in 150 μM biotin for 40 h before lysis and purification on streptavidin resin. Proteins were identified by mass spectrometry and each Nup-BioID sample was compared to the untagged parental strain negative control (Data S1).

To get an initial set of candidates, we filtered those proteins with a minimum 2-fold change over the parental control and ranked by the number of datasets in which the candidates were enriched. We further filtered these candidates based on predicted localization to other organelles based on potential targeting signals [33], domains functioning distinct from the NPC (*e.g.* transcription factors), and published subcellular fractionation data [34]. Finally, we closely examined predicted secondary structure and domains to identify patterns and sequence motifs that are often found in Nups (Figures 2-3). Encouragingly, this list included 13 of the previously predicted Nups, 7 of which had been previously validated. We added Nup43/TG*_310610 to our list of candidates, as it had been previously predicted as a Nup, but had been left unvalidated. This left us with 29 candidates to interrogate as potential Nups.

**Figure 2:**
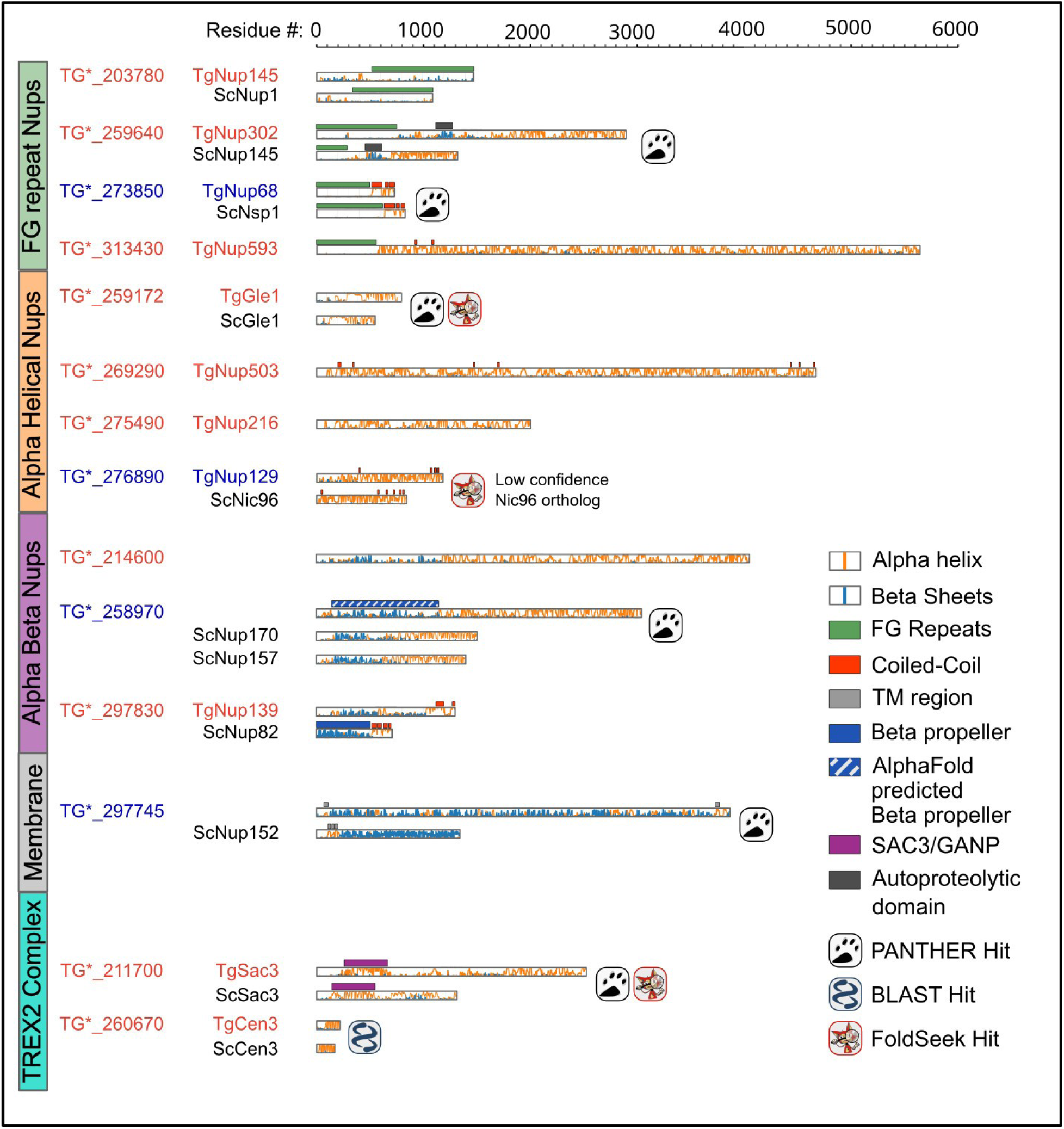
Domain architecture of essential and annotated Nups in Toxoplasma. The *Toxoplasma* Nups in red (essential Nups) or blue (non-essential) are divided into groups based on the Nup type, which are shown as vertical rectangular box on the left. The group type is adapted from [15,96]. Below the *Toxoplasma* Nups are the predicted yeast orthologs in black marked by the prediction type (PANTHER, BLAST, or Foldseek). The proteins are indicated by rectangular boxes where the protein length is in-scale to the residue numbers at the top. The secondary structure probabilities for helix (orange) and beta sheet (blue) are graphed inside the rectangular boxes. The boxed symbols in bottom right corner describe the predicted domains marked above the protein boxes.

**Figure 3:**
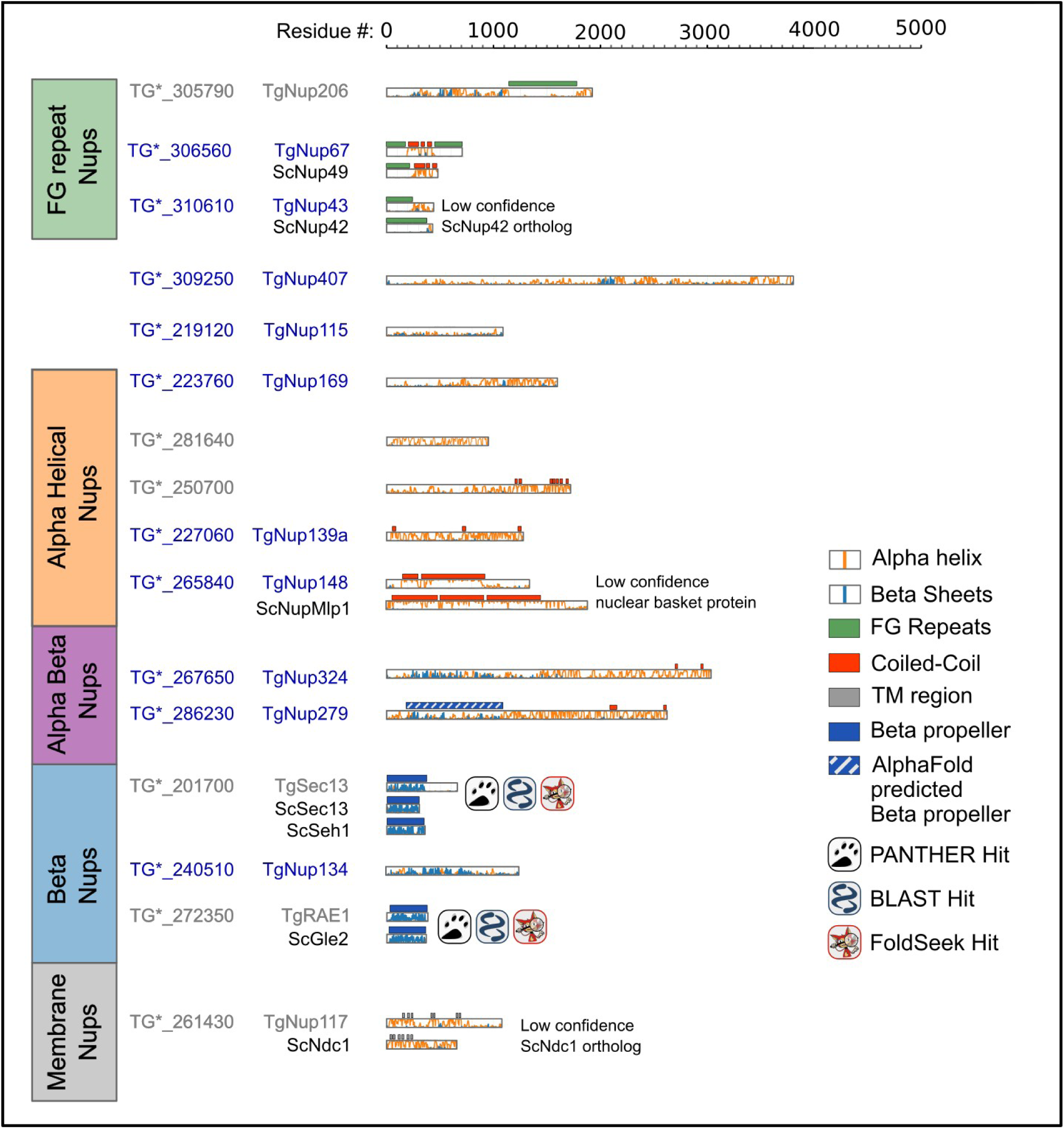
Domain architecture of non-essential and unverified Nups in Toxoplasma. *Toxoplasma* Nups in blue (non-essential Nups) or gray (unvalidated) are divided into groups as shown in Figure 2. All annotations are as in Figure 2.

We next sought to validate the localization of these proteins to the NPC. Of the 29 potential Nups, we successfully generated 23 strains in which the candidates expressed a C-terminal tag comprising an AID degron [35] and 3xHA epitope tag in the background of a strain in which the Nup302C protein was fused to a C-terminal 6xTy1 epitope. We confirmed correct integration of the AID-3xHA tag in these strains by PCR amplification (Figure S1). We then used confocal microscopy to visualize the subcellular localization of each protein. We found that 15 of our 23 tagged candidates localized to the NPC (Figures 2, S2), and were therefore bona fide Nups. Notably, we observed no convincing NPC localization for 1 of the previously predicted Nups (“Nup37”/TG*_248500 [20]; Data S2), in spite of its strong homology to yeast Nup57.

### The Toxoplasma nuclear pore complex is robust to nucleoporin loss

The NPC is an enormous protein complex that carries out an essential cellular function. Nevertheless, the NPC appears remarkably robust to loss of individual components. For instance, in yeast, only 12 of 33 Nups are essential [36]. To test the fitness costs associated with loss of the Nups we identified above, we used the auxin-inducible degron system. We first used immunofluorescence microscopy to confirm the knockdown of each Nup^AID^ protein after growth of parasites in media with the auxin indole-3-acetic acid (IAA). To test for fitness costs, parasites were plated on fibroblast monolayers in ±IAA media, and allowed to grow for 7-10 days. This allows for multiple rounds of invasion, growth, and lysis to occur, producing plaques where the monolayer has been disrupted. We tested the fitness cost of the 15 Nups we had validated, as well as Nup68 and Nup134, which have not been previously phenotypically characterized. We found that 7 of the Nups were essential (Nup145/TG*_203780, Sac3/TG*_211700, Nup433/TG*_214600, Nup218/TG*_226950, Nup216/TG*_275490, Nup139b/TG*_297830, Nup593/TG*_313430); induced degradation of these proteins resulted in a complete loss of plaque formation (Figure 5A, S5). We also found that degradation of an additional 4 Nups (Nup169/TG*_223760, Nup148/TG*_265840, Nup413/TG*_297745, Nup43/TG*_310610) resulted in significant fitness cost, though the proteins were not essential (Figure 4B,5D).

**Figure 4:**
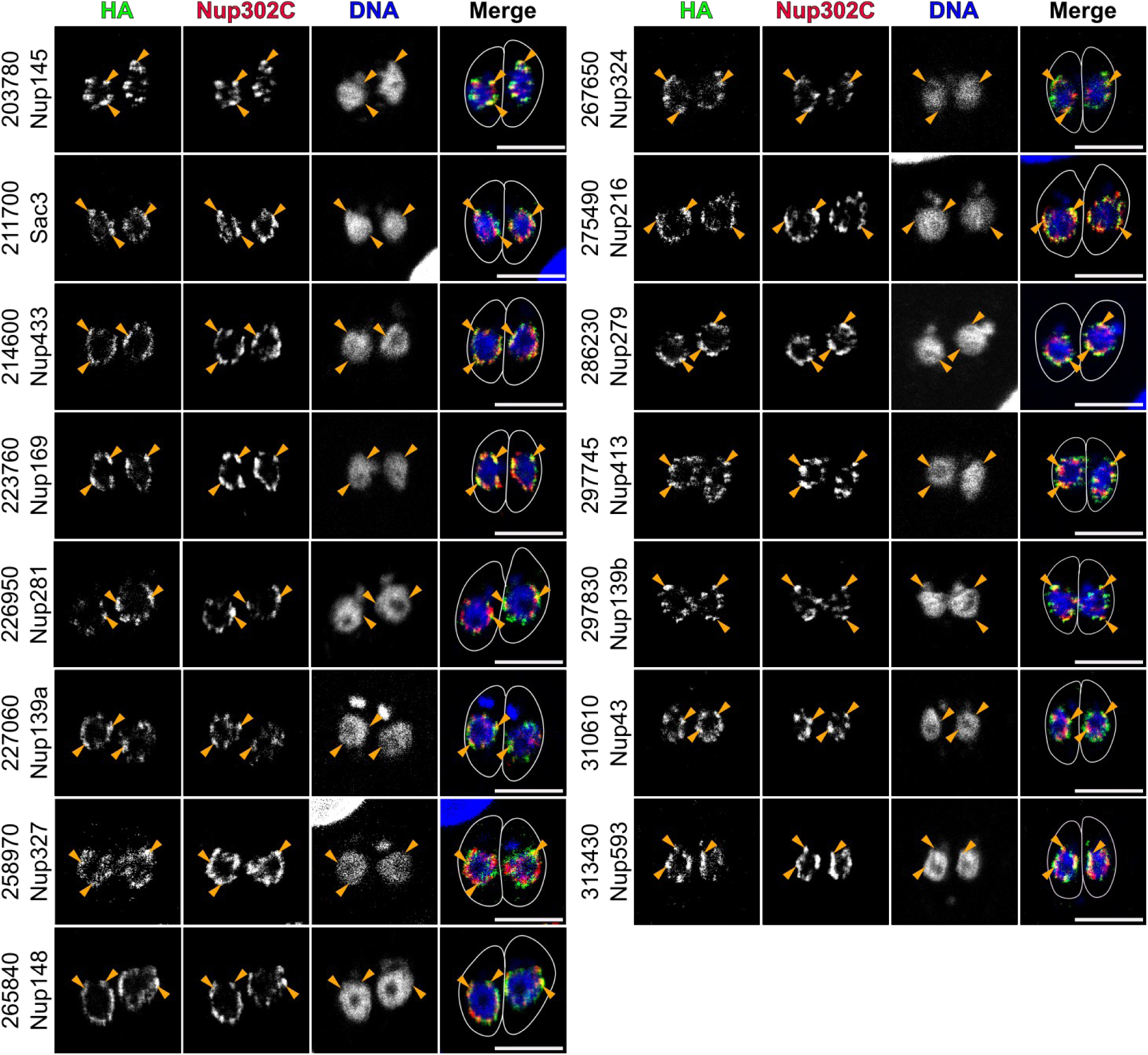
Validation of NPC localization for candidate Nups. Intracellular parasites with the indicated AID-3xHA tagged candidates were stained with antibodies recognizing HA (green), Ty1 (red; Nup302C), and Toxoplasma β-tubulin (not shown; used to create outline). DNA was stained with Hoechst (blue). Images are 1 airy-unit confocal slices (∼0.75 μm). Clear NPC-associated staining is noted with orange arrowheads. All scale bars are 5 μm.

**Figure 5:**
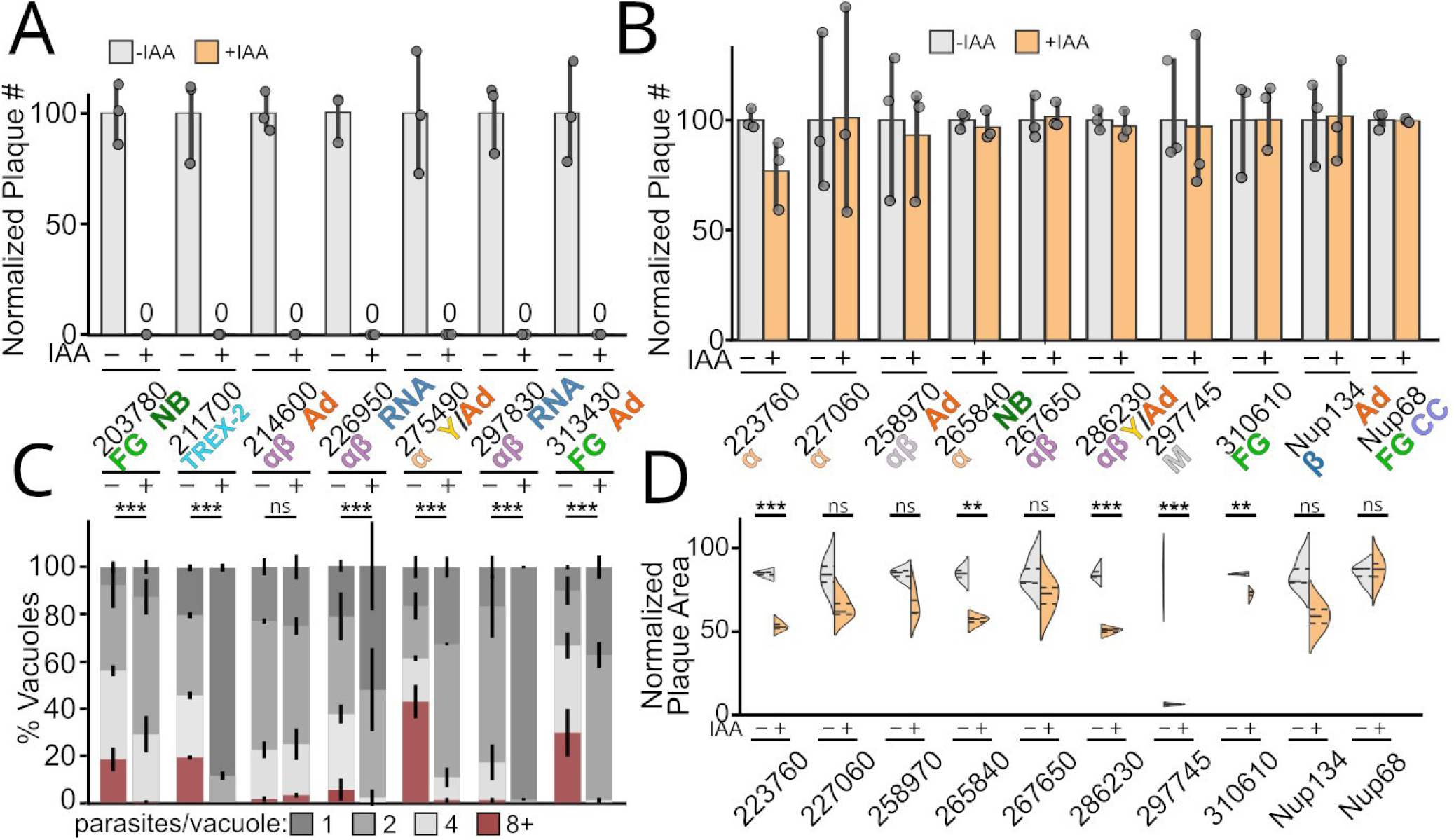
The *Toxoplasma* NPC is robust to loss of individual proteins. (A) Growth in +IAA media completely blocked the ability of 6 Nups to form plaques on a fibroblast monolayer. (B) The same strains were grown for 18 h post-infection ±IAA and showed varying levels of block in replication (see also Figure S6). ***, p<0.001 by Kolmogorov-Smirnof test. Quantification of 100 vacuoles from per n=3 biological replicate. (C) Other Nups showed no significant loss in plaque efficiency when grown in +IAA media. (D) Quantification of plaque area indicates a fitness cost upon depletion of a subset of these non-essential Nups. n=3 biological replicates. **, p<0.01; *** p<0.001 by unpaired two-tailed Student’s t-test. Nup groups are indicated as described in Figures 2-3 (α-helical, orange; β-sheet, blue; α-helix/β-sheet, purple; FG-repeat, green; Membrane, gray).

The reduction in plaque number seen in the case of essential Nups can be due to a block in replication, or motility and invasion. Some Nups, such as human Nup62 (TgNup68), have additional localization associated with functions independent of their roles in nucleocytoplasmic transport [37,38]. In addition, some FG repeat Nups have recently been implicated to be moonlighting as barrier-forming proteins at the base of the cilium in animals [39–41]. The *Toxoplasma* invasion machinery is thought to have evolved from the eukaryotic cilium [42]. We therefore sought to determine whether the essentiality of the above *Toxoplasma* Nups was due to a block in replication or at another step in the lytic cycle. We grew each of the 7 essential Nup^AID^ strains in ±IAA media for 16 hours and quantified the number of parasites per vacuole (Figure 5C). We found that loss of 2 of the Nups (Sac3/TG*_211700 and Nup139b/TG*_297830) caused a severe block in replication, in which the majority of parasites failed to divide even once. 4 of the validated Nups (Nup145/TG*_203780, Nup218/TG*_226950, Nup216/TG*_275490, Nup593/TG*_313430) showed significant defects in replication at 16 h, and exhibited aberrant morphologies after 24 h growth in +IAA, demonstrating that they were nonviable (Figure 5C,S5). Induced degradation of the last Nup, Nup433/TG*_214600, showed no significant effect on replication after 16 h growth in +IAA. However, these parasites also failed to replicate past 24 h in +IAA medium, again suggesting nonviability (Figure S6B). Taken together, these data are consistent with essential roles in NPC function rather than any specialized moonlighting function in invasion or motility.

### Ultrastructure expansion microscopy reveals relative positions of essential Toxoplasma Nups in the NPC

We used ultrastructure expansion microscopy (UExM) [43] to gain a higher resolution view of our newly identified essential Nups. These images allowed us to determine the relative positions of these proteins in the *Toxoplasma* NPC (Figure 6,S7). As above, we used Nup302C^6xTy1^ as a control, a known component of the Y-shaped complex, which is restricted to the nucleoplasmic side of the NPC in *Toxoplasma [7]*. As expected, all of the Nups we tested form well-distinguished puncta around the nucleus proximal to Nup302C (Figure 6,S7). Consistent with their predicted positions in the central ring (Figure 1A) Nup68 and Nup134 were respectively localized 34±6 nm and 45±7 nm towards the cytoplasmic side of the NPC from Nup302C, Intriguingly, Nup218/TG*_226950 and Nup139b/TG*_297830 appear to be on the extreme cytoplasmic face of the NPC, as they are even further from Nup302C, at 75±8 and 83±6 nm, respectively. Nup433/TG*_214600 (22±9 nm) and Nup593/TG*_313430 (47±5 nm) are both located a similar distance from Nup302C as Nup68 and Nup134, suggesting they are also adapter Nups or part of the central channel. Nup216/TG*_275490 appears to lie somewhere between Nup302C and the central channel, as it is only 11±6 nm towards the cytoplasm from Nup302C. Nup145/TG*_203780 colocalizes with Nup302C (6±10 nm), suggesting it is also a component of the Y-shaped complex. Finally, Sac3/TG*_211700 localizes 31±5 nm towards the nucleoplasm from Nup302C. Thus, the essential Nups are well distributed across the NPC. Nup148/TG*_265840 (previously called nuclear mesh protein 1 or NMP1 [44]), while non-essential, has a predicted extended coiled-coil domain similar to that present in the the yeast and human nuclear basket spoke proteins (Figure 3). Because no density was observed for the *Toxoplasma* nuclear basket by cryo-ET, we sought to test whether Nup148/TG*_265840 was located at a position consistent with a potential role in the nuclear basket. UExM demonstrated Nup148/TG*_265840 is a definitive Nup with puncta that colocalize closely with Nup302C (2 ± 14 nm; Figure S7). Thus Nup148/TG*_265840 localizes to the nuclear side of the NPC, and may be part of the *Toxoplasma* nuclear basket.

**Figure 6:**
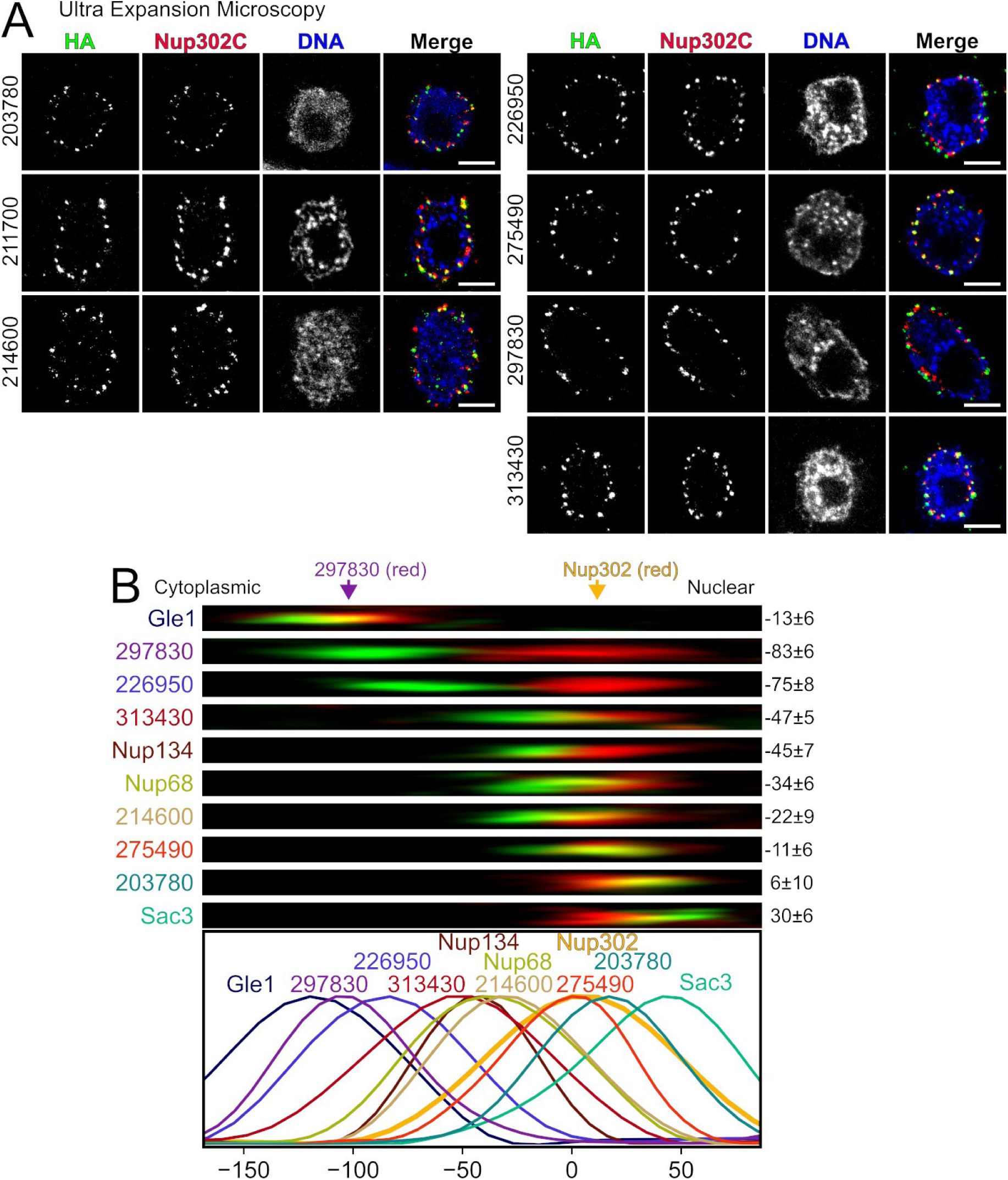
UExM reveals relative position of individual Nups in the NPC. (A) UExM images are maximum intensity projections of 10 slices from the middle of the parasite nucleus for essential candidate Nups. All scale bars are 5 μm (∼5× expanded, ∼1 μm in actual size). (B) Representative line scan profiles used to measure distance between indicated candidate Nup (green) and NPC marker (Nup302C or Nup139b/Nup297830 – Gle1 only). Representative images for each line profile are above the plots. Mean distances with sem are indicated to the right of each image (minimum n=10 puncta). Additional UExM images in Figure S7.

### The Toxoplasma Nups TG*_211700 and TG*_297830 are required for mRNA export from the nucleus

One of our candidates had a predicted role in mRNA export. TG*_211700 has significant homology to Sac3, which forms the central scaffold of the TREX-2 nucleoplasmic mRNA export machinery (PDB: 4MBE and 8U8D; PTHR12436 – e-value: 4×10^-37^; Data S2) [45,46]. Consistent with this idea, we found Sac3/TG*_211700 localizes at the extreme nucleoplasmic face of the NPC when imaged by UExM (Figure 6). We tested whether induced degradation of Sac3/TG*_211700 and the other essential nucleoporins affected nuclear mRNA export in *Toxoplasma*, as mRNA export is an essential function of the NPC, the disruption of which would be expected to cause an immediate block in replication. We expected mRNA export to be disrupted immediately upon loss of the proteins involved (Figure 7). We found that 2 h incubation with IAA was sufficient for complete degradation of each of the essential Nups (Figure S3), so chose this as our experimental timepoint. Parasites were allowed to infect host cells in normal media for 16 h, and then changed to ±IAA media for another 2 h before fixation. Mature mRNA was stained by fluorescent *in situ* hybridization (FISH) using a fluorescein poly(dT) oligonucleotide. Parasites were counterstained with Hoecsht, and the nuclear poly(dT) signal for each parasite was quantified. Consistent with a predicted role in mRNA export, Sac3/TG*_211700 showed a significant increase in mRNA build up in the nucleus upon growth in +IAA (Figure 7A,B). Only one other candidate, Nup139b/TG*_297830, also showed a clear block in mRNA export upon growth in +IAA. This protein, like Sac3/TG*_211700, also showed the fastest-onset replication defect we observed (Figure 5). In contrast to Sac3/TG*_211700, Nup139b/TG*_297830 localizes to the cytoplasmic side of the NPC (Figure 6). The secondary structure profile of TG*_297830 is similar to that of yeast ScNup82 (Figure 2). ScNup82 forms the core of the cytoplasmic mRNA export complex of the yeast NPC and, like Nup139b/TG*_297830, is essential for mRNA export [47]. While the two proteins have no detectable primary sequence homology, these observations strongly argue that Nup139b/TG*_297830 is the *Toxoplasma* ortholog of ScNup82, or at least serves an analogous function.

**Figure 7:**
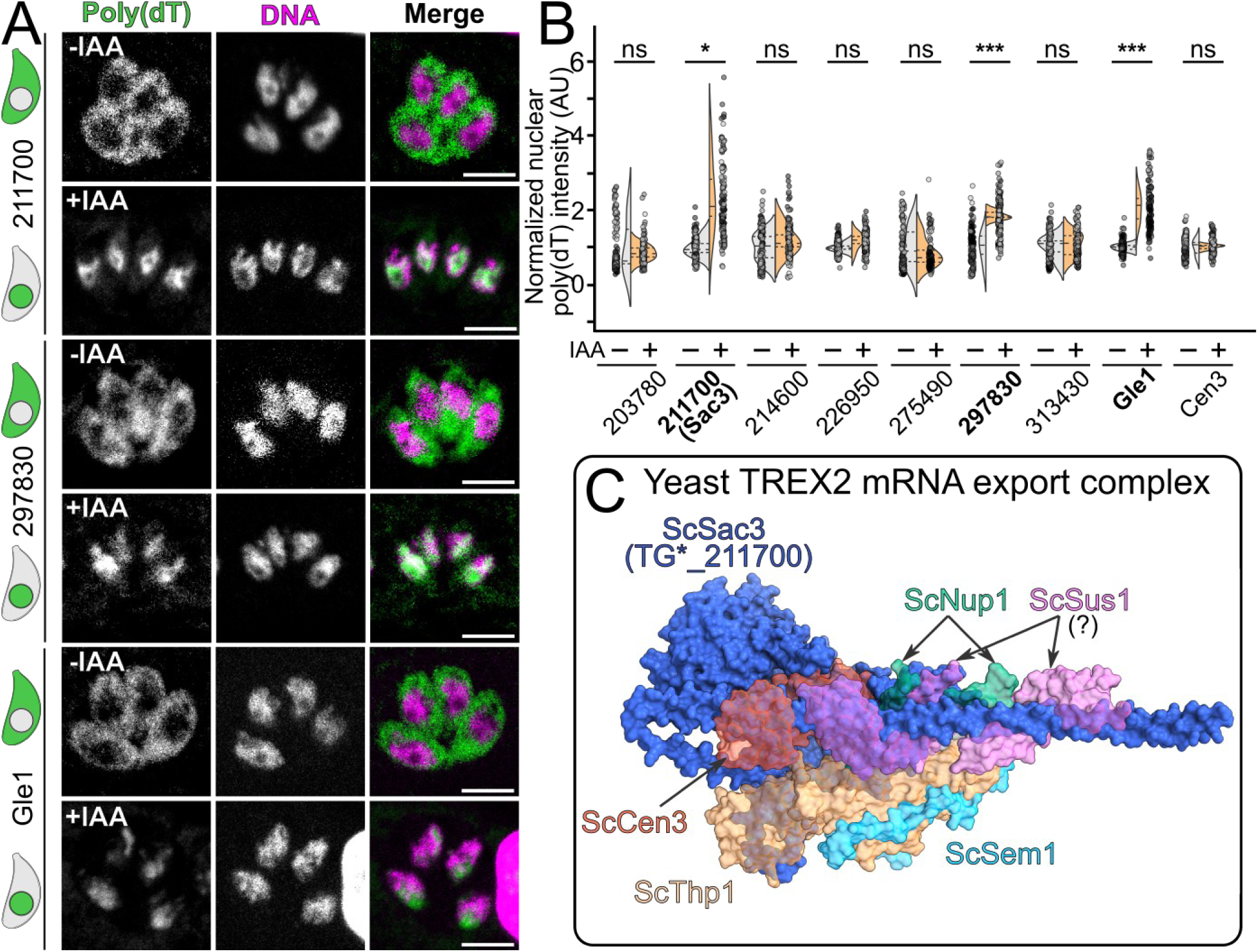
Identification of *Toxoplasma* Nups required for mRNA export. (A) Confocal slices of parasites that were allowed to infect overnight and then grown in +IAA media for 2 h before fixation. mRNA was stained by FISH with poly(dT) oligo (green) and parasites were counterstained with Hoechst (magenta). Scale 5 µm. Cartoon represents a healthy parasites next to -IAA and a parasite with mRNA export block next to +IAA. (B) Quantification of images from (A). A minimum n=20 vacuoles were quantified per n=6 biological replicates. Individual points are represented as circles with replicates colored in different shades of gray. ns, not significant; ***, p<0.001; two-tailed unpaired Student’s t-test. (C) Model of the yeast mRNA export complex demonstrates well-folded domains for each of the components. Orthologs of ScNup1 and ScCen3 appear present in *Toxoplasma* but appear uninvolved in mRNA export. No clear Sus1 ortholog is present in *Toxoplasma*.

Our bioinformatic, localization, and FISH data suggest that TG*_211700 is, indeed, a functional ortholog of Sac3. Unlike many proteins associated with the NPC, the TREX-2 complex is composed of proteins that have well-defined globular folds (Figure 7C). We reasoned this should facilitate prediction of other members of the TREX-2 complex in *Toxoplasma*. Using HMMER, we identified TG*_236920 to be a likely ortholog Sem1. Sem1 is a component of the proteasome that also serve in the TREX-2 complex [48] (Figure 7C,S8-9). While we identified potential orthologs of Sus1/ENY2 in Chromerids, ciliates, and dinoflagellate lineages, this protein appears to be missing throughout Apicomplexa (Figure 7C,S10,Data S3-4). Thp1 is another member of the TREX-2 complex, and is defined by its PCI-domain. HMMER identified TG*_264060 as the closest match to a Thp1-like profile (PTHR12732:SF8), though our phylogenetic analysis was inconclusive as it did not confidently cluster known Thp1 orthologs away from paralogs with other functions. Unfortunately, we were unable to tag and thereby validate Sem1/TG*_236920 or Thp1/TG*_264060. We did, however, identify *Toxoplasma* orthologs of other components of mRNA export machinery. Centrin-3, also called Cdc31 in yeast, helps anchor the TREX-2 complex in other organisms [49,50]. Centrin-3 has a previously described ortholog in *Toxoplasma* [51]. We therefore created a strain in which TgCentrin-3 (Cen3) had been tagged with AID-3xHA, as with our other candidates (Figure S7). We tested this strain for a defect in mRNA export, but saw no significant difference between ±IAA conditions (Figure 5), suggesting that, unlike yeast Cen3/cdc31 [50], TgCen3 is not a central player in this process. We identified TG*_259172 as sharing significant homology with yeast Gle1 (PTHR12960 – evalue: 3×10^-37^; Data S2-S3), a protein essential for both mRNA export and translation termination in other organisms [52,53]. Consistent with this observation, when we created a TgGle1^AID-3xHA^ strain, we found that it localized to the cytoplasmic side of the NPC (Figure 6B,S7) and induced degradation of the protein caused a striking buildup of mRNA in the nucleus (Figure 7A,S11) explaining the essentiality of Gle1 for parasite survival (Figure S11B). These data indicate that TgGle1 is indeed an ortholog of the yeast protein and has critical role in mRNA export in *Toxoplasma*. Together our data highlight that, like the NPC itself, the mRNA export machinery in *Toxoplasma* has conserved central components, but has diverged in its specific architecture.

### Identification of Nups required for NPC stability or assembly

Another way in which a nucleoporin might be essential is by stabilizing existing NPCs or by facilitating the biogenesis of new NPCs. We therefore tested whether inducing degradation of any of the 7 essential Nups caused mislocalization of our established NPC marker, Nup302C^6xTy1^, which is part of the Y-shaped complex on the nucleoplasmic face of the NPC (Figure 8A). We also tested for mislocalization of Nup139b/TG*_297830, as our data demonstrate it is on the opposite side of the NPC (Figure 6). To this end, we created parasites in which the endogenous Nup139b/TG*_297830 protein was expressed as a C-terminal fusion to 6xTy1 and tagged the other 6 essential Nups with AID-3xHA in this background. Parasites were allowed to infect host cells for 2 h in normal media and then switched to ±IAA media for another 8 h before fixation. In all cells grown in -IAA, both marker Nups showed characteristic punctate staining on the nuclear envelope (Figure 8B,S12-13). Knockdown of 6 of the 7 Nups caused aggregation and mislocalization of Nup302C. Similarly, loss of 4 of the 6 Nups caused aggregation and mislocalization of Nup139/TG*_297830 (Figure 8B,S12-13).

**Figure 8:**
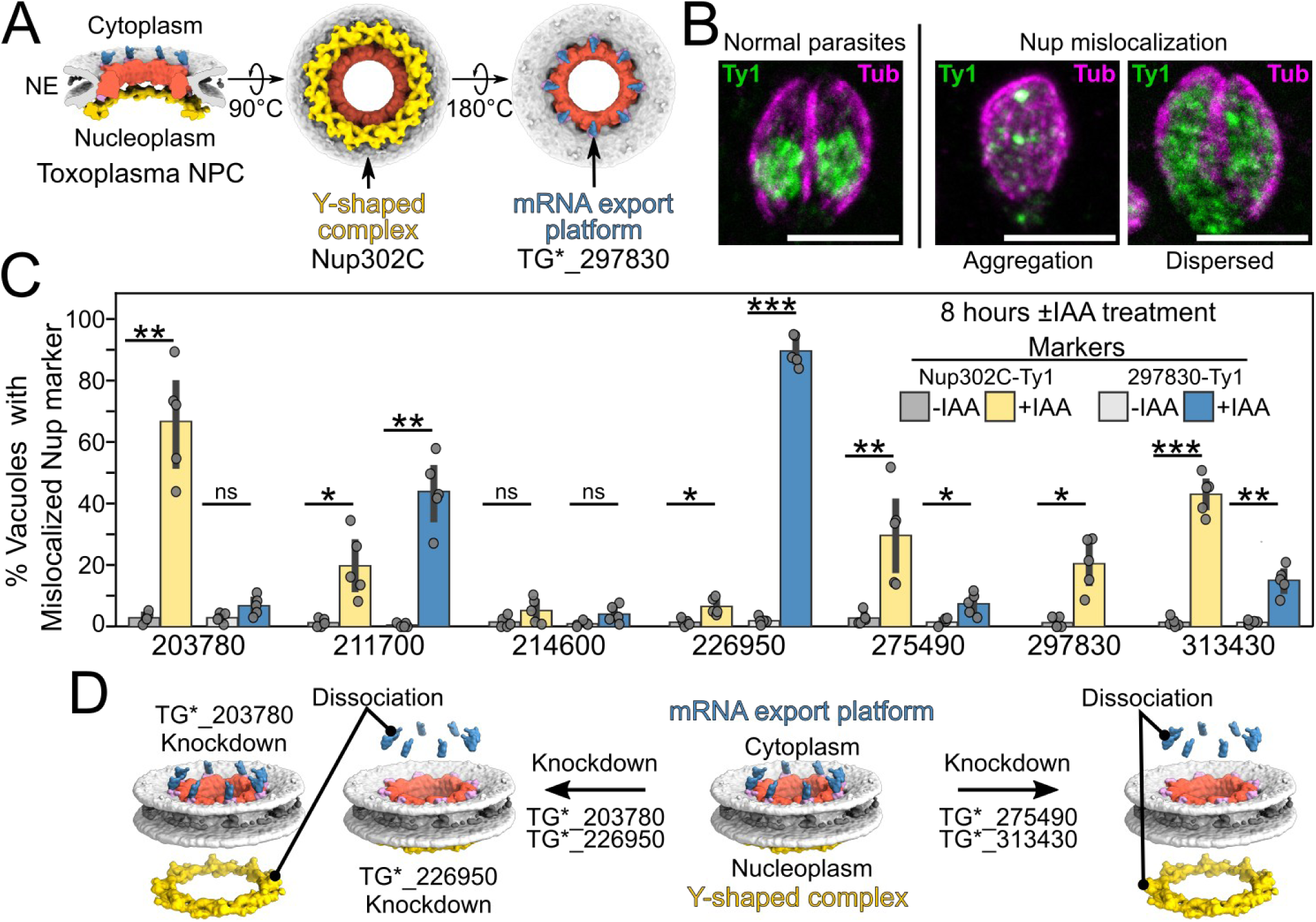
Identification of Nups required for assembly and stability of NPC. (A) Model of the *Toxoplasma* NPC from [7] indicates likely localization of Nup302C in the nucleoplasmic Y-shaped complex (yellow) and Nup139b/TG*_297830 in the cytoplasmic mRNA export platform (blue). (B) Parasites showing normal and mislocalized NPC marker (green). Images are maximum intensity projection of confocal slices. Scale 5µm. More examples shown in Figures S12, S13. Note nuclear punctate staining of marker Nups (green) centered in tubulin outline (magenta). (C) Quantification of mislocalization of the indicated marker in parasites that had been grown for 8 h ±IAA post-infection. Note that because Nup139b/TG*_297830 is used as a marker, we quantified only mislocalization of Nup302C. (D) Cartoon summary of data indicating knockdown of certain Nups affects dissociation of specific subcomplexes from the *Toxoplasma* NPC.

Consistent with the rapid block of replication upon their loss, knockdown of either Nup139b/TG*_297830 or Sac3/TG*_211700 showed a modest penetrance of the mislocalization phenotypes (Figure 8C). Given the substantial effects of their loss on Nup localization at 8 h (Figure 8C), we sought to ascertain which phenotype associated with these proteins was primary (mRNA export or NPC stability). To this end, we tested whether loss of either protein caused a significant defect in Nup marker localization over the same short 2 h time period we used for the mRNA export assay. We also tested whether loss of Nup139b/TG*_297830 affected Sac3/TG*_211700 localization to the NPC at this time point. For all markers tested, loss of either protein caused no significant disruption to the NPC architecture in this timeframe (Figure S11C). Our data therefore suggest that both Sac3/TG*_211700 and Nup139b/TG*_297830 directly function in the export of mRNA in *Toxoplasma* and the severe mislocalization of markers at later time points is a secondary effect of this function.

Loss of Nup216/TG*_275490 and Nup593/TG*_313430 also caused mislocalization of both markers, though with a modest penetrance of only 5-40%. Given their colocalization with the central channel proteins Nup68 and Nup134 by UExM (Figure 6), we reason that these two proteins are also part of central adapter complex, which is consistent with our predictions based on secondary structure (Figure 2-3, Data S2). Notably, degradation of Nup281/TG*_226950 caused an almost complete dispersion of the cytoplasmic-facing Nup139b/TG*_297830 into the cytoplasm in >90% of cells, but only a mild mislocalization of nucleoplasmic Nup302C in <10% of cells (Figures 8C,S12-S13). Nup281/TG*_226950, like Nup139b/TG*_297830, lies on the cytoplasmic side of the NPC. To know if this specific effect is because Nup281/TG*_226950 is required to anchor Nup139b/TG*_297830 stably on the NPC, we quantified Nup139b/TG*_297830 mislocalization at earlier timepoints after 2 h and 4 h of IAA treatment. Consistent with a role in the stability, rather than assembly of Nup139b/TG*_297830 on the NPC, we see that Nup139b/TG*_297830 localization was completely dispersed at 2 hours of IAA treatment in the Nup281/TG*_226950^AID^ strain (Figure 9A,B). Taken together, our data suggest that Nup281/TG*_226950 is either in the same subcomplex as Nup139b/TG*_297830 or required for its stability on the NPC. In contrast, degradation of TG*_203780 resulted in the specific mislocalization of nucleoplasmic-facing Nup302C, but not TG*_297830 (Figure 8C,S12-13). We then tested if Nup145/TG*_203780 knockdown affected Nup302C stability early by quantifying Nup302C mislocalization at earlier timepoints from 2-6 h (Figure 7A,C). We see that there is progressive increase in the mislocalization of Nup302C over time after Nup145/TG*_203780 knockdown (Figure 9A,C). Additionally, we did not see that the mislocalization was correlated with parasite division, but rather with increased time in +IAA media (Figure 9C). Notably, yeast ScNup1 is an FG repeat protein that is part of the nuclear basket and helps anchor the Y-shaped complex to the nuclear membrane and the rest of the NPC in other organisms (Figure 9D). Nup145/TG*_203780 is also an FG repeat protein and colocalizes with Nup302 in our UExM imaging (Figure 6). Together, these data strongly suggest Nup145/TG*_203780 is a functional ortholog of (or functionally analogous to) ScNup1 and a potential component of the *Toxoplasma* nuclear basket.

**Figure 9:**
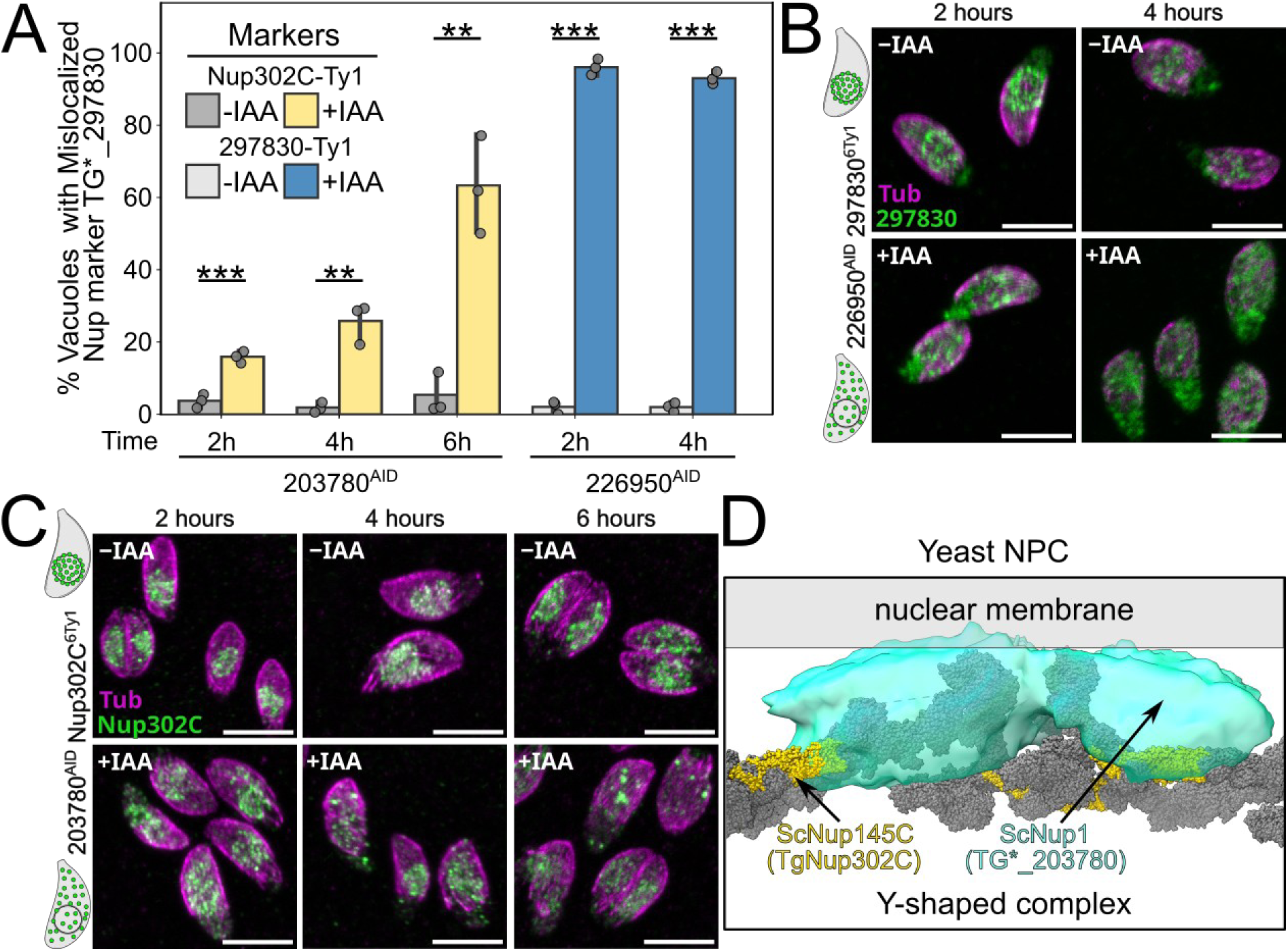
Nup145/TG*_203780 and Nup281/TG*_226950 are required for stability of specific subcomplexes. (A) Quantification of NPC marker mislocalization in the indicated strains that had been grown for 2-6 h in ±IAA after infectoin. ***, p<0.0001; two-tailed unpaired Student’s t-test. Maximum intensity projections of confocal images showing (B) mislocalized Nup139b/TG*_297830^6xTy1^ upon knockdown of Nup281/TG*_226950 and (C) mislocalized Nup302C^6xTy1^ upon knockdown of Nup145/TG*_203780. Scale bars are 5 µm. Cartoon guide of observed staining next to each set of images. (D) CryoEM model of the yeast Y-shaped complex shows likely localization of Nup302C (yellow) and ScNup1 (cyan volume), which helps anchor the Y-shaped complex.

The critical function of the NPC is mediating nucleocytoplasmic transport of proteins and mRNA [54]. We had already identified Sac3/TG*_211700 and Nup139b/TG*_297830 and as essential for mRNA export and Nup281/TG*_226950 and Nup145/TG*_203780 as important for maintaining NPC stability. Any effect on NCT by degradation of these proteins is likely to be secondary. We therefore focused on determining whether knockdown of either Nup216/TG*_275490, Nup593/TG*_313430, or Nup433/TG*_214600 caused a measurable loss in protein NCT. To this end, we created an NCT reporter strain by ectopically overexpressing the small nuclear protein TG*_258850 (identified in this study; Figure S2A) with a 3x-FLAG tag. We then tagged the remaining 3 essential Nups with AID-3xHA in this background. As a positive control, we used an NCT reporter strain with *Toxoplasma* RanGAP (TBC9) [55] tagged with AID-3xHA. Parasites were allowed to infect the host cells for 2 hours and then treated with ±IAA for 8 hours (Nups) or 2 hours (TBC9). As expected, knockdown of TBC9 caused immediate release of TG*_258850^3xFLAG^ from the nucleus to cytoplasm indicating a strong NCT defect (Figure 10), followed shortly thereafter by parasite death [55]. Perhaps unsurprisingly given the progressive phenotypes we observed for these strains (Figures 5,8), we found that loss of any of the 3 Nups tested had no significant effect on NCT in these parasites (Figure 8B). However, we found a small percentage of parasites in which the NCT reporter had begun to leak into the cytoplasm after Nup593/TG*_313430 knockdown (Figure 10A), which were never observed in the -IAA condition. Notably these parasites had recently completed division, suggesting Nup593/TG*_313430 is required for assembly of fully functional NPCs. This observation is consistent with our finding that Nup593/TG*_313430^AID^ parasites are able to replicate no more than once in +IAA media (Figure 5C).

**Figure 10:**
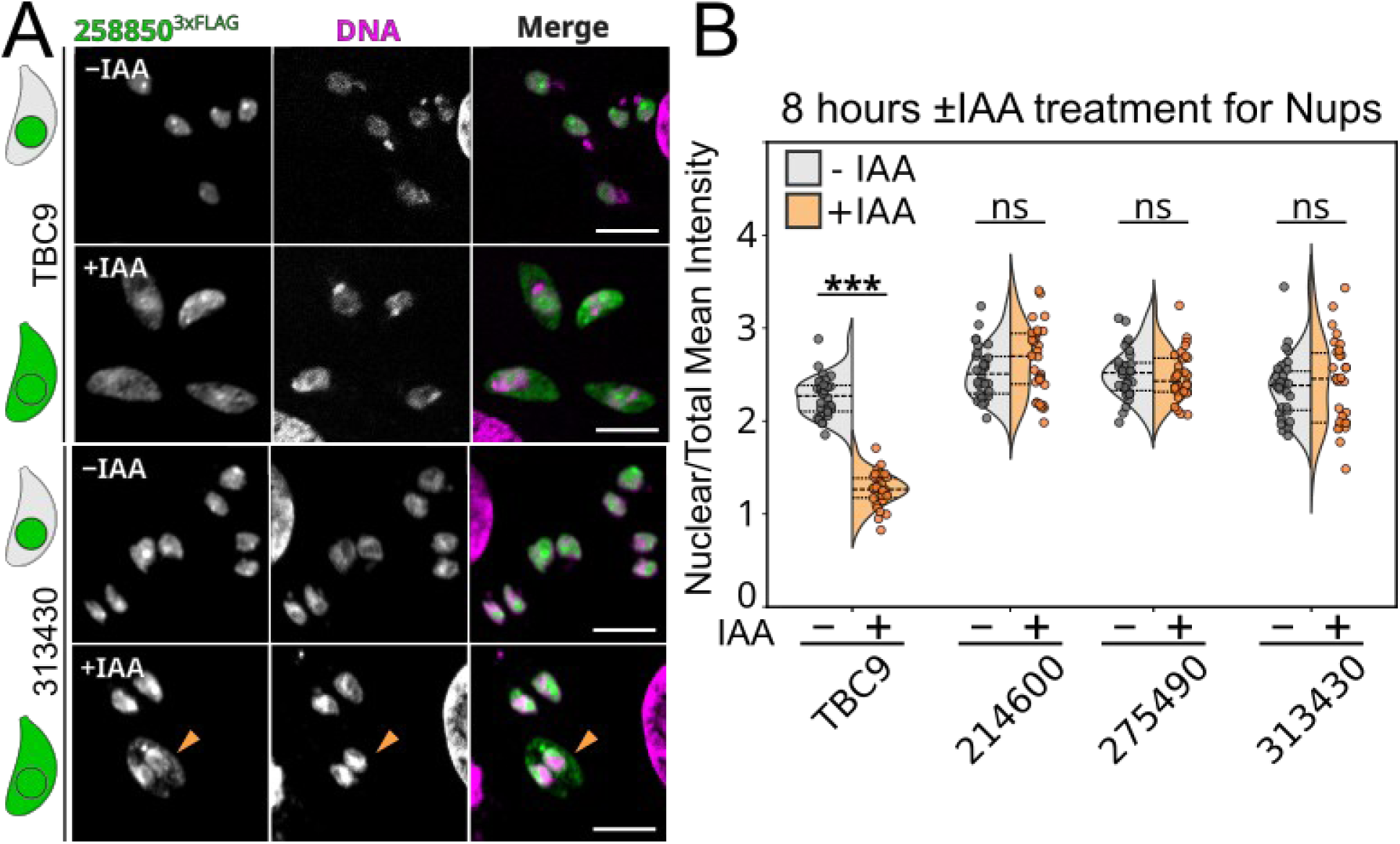
Nucleocytoplasmic transport is robust to the knockdown of individual Nups. (A) Confocal slices of a TBC9^AID^ (positive control) and TG*_313430^AID^ parasites expressing NCT marker TG*_258850^3xFLAG^ grown in ±IAA. A small population of TG*_313430^AID^ parasites have a modest NCT defect (orange arrowhead). Note these parasites have recently divided. Scale bars are 10 µm. Cartoon guide of expected phenotype staining (loss of nuclear retention). (B) Quantification of the nuclear over total mean intensity for each condition. n=10 cells were averaged per n=3 biological replicates. ns, not significant; ***, p<0.0001; two-tailed unpaired Student’s t-test. Nup^AID^ were treated for 8 h in ±IAA, TBC9^AID^ for 2 h.

In addition to directly altering NPC stability and function, some Nups are required for efficient assembly of new NPCs [56,57]. Because UExM revealed NPCs as individual puncta (Figure 6), we reasoned we could use this technique to accurately quantify NPC number on the *Toxoplasma* nucleus (Figure 11). Because an assembly defect is by its nature progressive, we chose the possible latest timepoint of +IAA treatment for each strain. Since Nup593/TG*_313430^AID^ parasites are unable to undergo a second replication event after 12-16 in +IAA, we treated them with IAA for 12 hours to ensure the majority of parasites had the opportunity to divide once. Nup216/TG*_275490^AID^ and Nup139b/TG*_214600^AID^ parasites are unable to replicate after ∼24 h in +IAA media, and were therefore treated with IAA for 18 hours. We included the Nup139b/TG*_297830^6xTy1^ parental strain as a negative control. Because of the mislocalization of aggregated species upon treatment with +IAA (Figure 8), we considered only those puncta localized to the periphery of DNA staining as bona fide NPCs (Figure 11). In the parental strain, TG*_297830^6xTy1^, we counted 56±3 puncta per nucleus, which was unaffected by treatment with IAA. Surprisingly, knockdown of Nup593/TG*_313430 did not show a significant defect in assembly (51±6 versus 49±5 puncta in ±IAA). However, knockdown of Nup433/TG*_214600 produced a dramatic reduction from 70±5 to 47±4 puncta per nucleus. Notably, we observed that Nup433/TG*_214600^AID^ nuclei appear to have a higher number of NPCs than the parental strain when grown in -IAA. This suggests some compensatory effect in these cells due to a fitness cost of tagging Nup433/TG*_214600, perhaps due to non-specific “baseline” degradation through the AID system [58,59]. While we counted 59±4 puncta on the nuclei of Nup216/TG*_275490^AID^ parasites grown in -IAA and 47±4 puncta when grown in +IAA, the difference was not significant (p=0.07). However, we observed large foci on the periphery of the nucleus only in parasites grown in +IAA consistent with aggregation of NPCs upon knockdown of Nup216/TG*_275490 (Figure 11C). Of the three Nups we tested, only the predicted adapter protein Nup433/TG*_214600 appears to be required for assembly of new NPCs.

**Figure 11:**
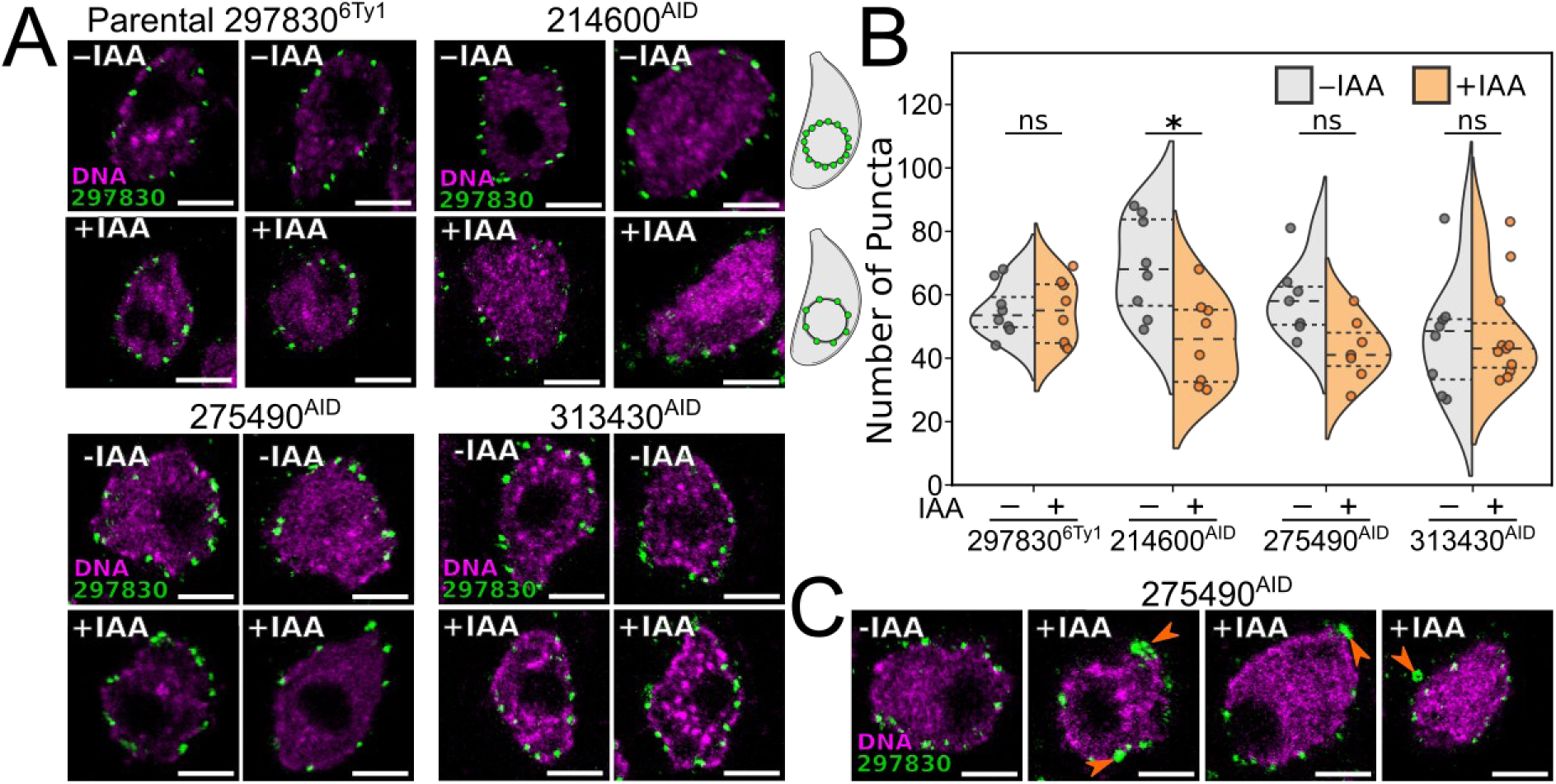
Loss of TG*_214600 causes reduction in NPC number. (A) Maximum intensity projections of representative UExM images for each strain characterized. Scale bars 5 µm (∼5× expansion). Cartoon parasites show reduction NPC expected for an assembly defect. (B) TG*_297830-positive NPC puncta were counted from parasites treated with auxin for 12 h (TG*_313430^AID^) or 18 h (TG*_214600^AID^ and TG*_275490^AID^). ns, not significant; *, p<0.01; Kolmogorov-Smirnov test (K-S test). (C) Maximum intensity projections of UExM images show aggregated NPC puncta on the nuclear envelope in the +IAA condition in TG*_275490^AID^ parasites.

### Most Toxoplasma Nups have no clear orthologs in model organisms

The individual protein components of the NPC appear to be remarkably widely conserved among eukaryotes, as clear orthologs to yeast proteins have been found in protists such as trypanosomes [15,25,60] and *Tetrahymena [24].* To gain an idea of the conservation of *Toxoplasma* Nups across taxa, we first used the OrthoMCL database [61,62]. We found that only 8 of the 29 (24 now validated and 5 still unvalidated) *Toxoplasma* Nups belonged to OrthoMCL groups including diverse taxa (Data S2). Instead the majority of *Toxoplasma* Nups belong to OrthoMCL groups with membership limited to Apicomplexa and, in some cases, the closely related Chromerids. To more deeply examine those proteins with well conserved domains, we used HMMER to identify related sequences from a broad sampling of eukaryotes from the EukProtv3 [63] database (Data S3-4). We then aligned these sequences and estimated phylogenetic trees to assess their evolutionary relationships.

This analysis confirmed that while Sec13 is broadly conserved in Alveolates (TG*_201700 in *Toxoplasma*) [27], its paralog Seh1 is found in ciliates but missing in other clades including dinoflagellates and Apicomplexa (Figure S14). Similarly, we confirmed that while the mRNA export protein Gle1 is conserved in Apicomplexa and dinoflagellates but missing in ciliates [24], both Sac3 and Gle2/Rae1 are broadly conserved in all alveolates (Data S3-4). One unusual case was Nup115/TG*_219120, which was validated as a Nup in a previous study [20]. This protein has a predicted zinc-finger domain that falls into the OrthoMCL OG7_0002161 group, which has members throughout Eukaryota. The human and yeast members are related to the RNA-binding protein ZFP36/TTP (also confusingly called NUP475 for “nuclear protein,” not to be confused with “Nup”), the founding member of a family of zinc-finger domains known to bind AU-rich RNA sequences and regulate mRNA decay [64,65]. ZFP36 is a nuclear-resident protein [64] with no known functional interactions with the NPC. Nup115/TG*_219120, on the other hand, localizes exclusively to the NPC [20]. We also identified two *Toxoplasma* paralogs of Nup115/TG*_219120, TG*_218300 and TG*_294840, which are likely more typical nuclear-localized RNA-binding proteins. We found that both paralogs appear broadly conserved in Apicomplexa and the closely related Chromerids, but not in more distantly-related alveolates such as dinoflagellates or ciliates (Figure S15 phylogeny). Together, these data suggest that Nup115/TG*_219210 may be an RNA-binding Nup that is a recent innovation in the Chromerid/Apicomplexa lineage.

We attempted to extend the prediction of orthology of each of the validated *Toxoplasma* Nups using a combination of primary sequence (BLAST [66]; the PANTHER domain database [67]) and structure-based (Foldseek [68]) methods (Figures 2-3, Data S2). Consistent with the OrthoMCL analyses described above, the strongest support by all methods are for proteins with well-folded domains, such as Sec13 and Gle2/Rae1 [20,27]. Two major motifs found in many Nups are β-propeller WD40 domains and the so-called FG repeats found in many of the proteins that line the pore of the NPC [69,70]. While many of the *Toxoplasma* Nups contain these motifs (Figures 2-3), their existence alone is not sufficient to predict orthologs. For instance, the only significant region of sequence similarity found by BLAST in most *Toxoplasma* FG repeat proteins are the FG repeats themselves, which are by their nature low complexity. Similarly, the predicted WD40 domains of many of the *Toxoplasma* Nups match to non-Nup proteins in other organisms when searched (Data S2). Foldseek searches are limited by the depth of sequence available to identify evolutionary relationships. We lack a robust set of sequences for organisms that link Apicomplexa to the last-eukaryotic ancestor, so AlphaFold models are often of much lower confidence than for other organisms. All together, we confidently identified orthologs to only 7 *Toxoplasma* Nup proteins (Reciprocal BLAST/Foldseek with e-value <10^-3^) Therefore, current tools for bioinformatic analyses appear ill-suited to inform on the potential functions of *Toxoplasma* Nups.

## Discussion

The *Toxoplasma* NPC has a distinct architecture from other model organisms [7], and nucleoporins that comprise this structure have been difficult to predict. Using a combination of proximity biotinylation and bioinformatic analysis, we have greatly expanded the functional catalog of *Toxoplasma* nucleoporins. Our work increased the number of proteins that have been experimentally validated as associated with the *Toxoplasma* NPC from 8 to 24. We saw no labeling of 2 previously validated Nups (Nup115/TG*_219120 and Nup407/TG*_309250) by proximity biotinylation. It is possible that these proteins are less accessible to biotinylation, either because they are not stable components of the NPC, or because of their localization in the NPC itself. The 6 candidates we were unable to endogenously tag include proteins with strong homology to known Nups with well-defined domains (Figure 2-3, Data S2), suggesting that these unvalidated proteins include additional bona fide Nups that merit future study. We have combined our predictions of domain architecture and potential homology (Figure 2-3, Data S2-4) with high-resolution UExM and phenotypic data to suggest subcomplex membership for each of our validated or well-predicted but as yet unvalidated Nups (Figure 12B). As the NPC in other organisms is comprised of ∼30 proteins [22,23], it is likely that our list now approaches the full complement of Nups in *Toxoplasma*.

**Figure 12:**
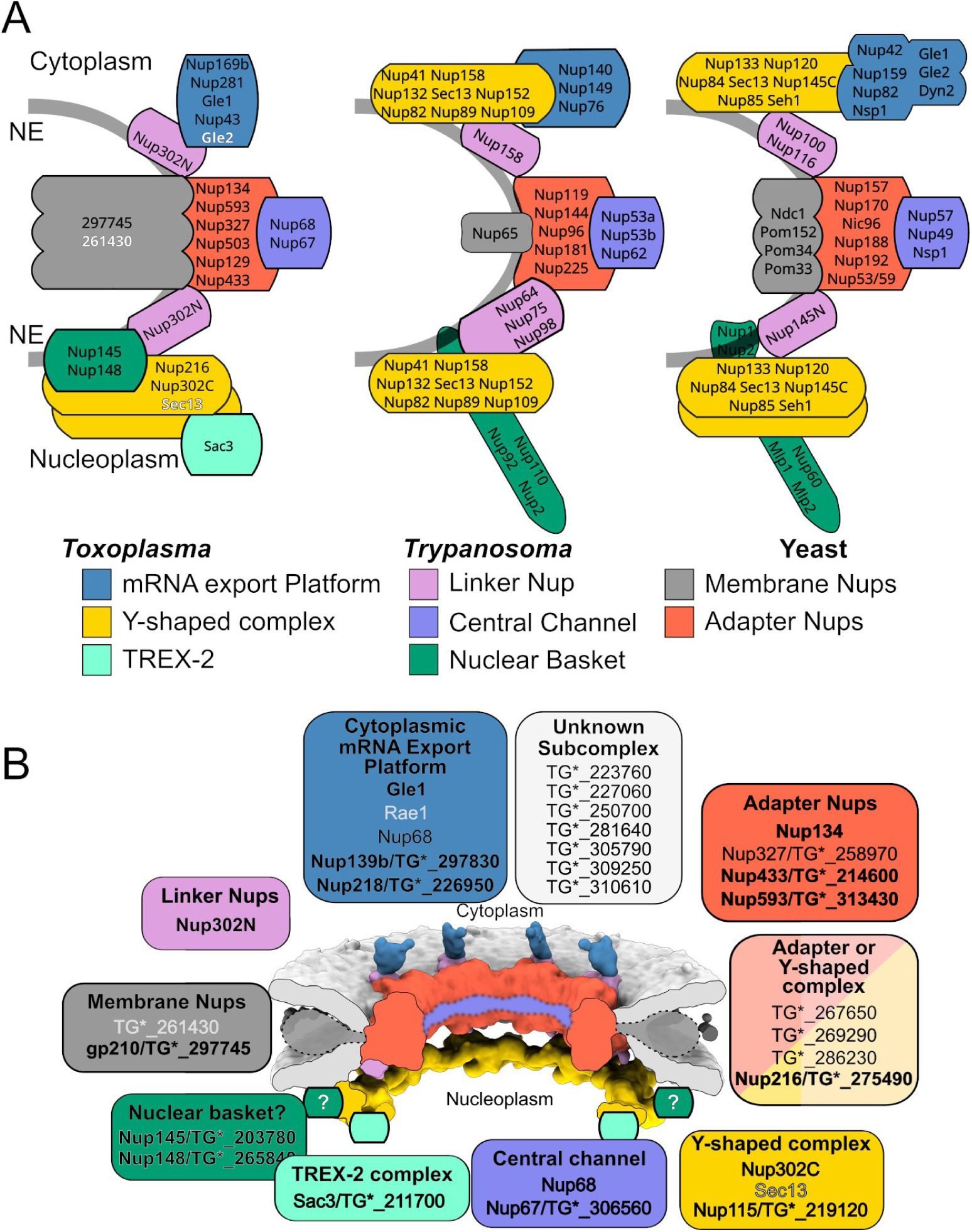
A comparative model of the *Toxoplasma* NPC subcomplexes. Cartoon representation of *Toxoplasma*, *Trypanosoma* [25], and Yeast [77] NPCs with their subcomplexes shown with the same color scheme as Figure 1. Nups are labeled inside their predicted or validated respective subcomplexes. (B) Model of the *Toxoplasma* NPC from cryoET maps [7] colored by subcomplex. Note that the densities for membrane Nups and central channel are poorly resolved, and have been replaced with cartoon representations. Nups have been grouped according to predicted subcomplex membership. Validated Nups in black text and bolded when we have high confidence of their membership in a given subcomplex, while predicted Nups that remain unvalidated are in gray text.

Surprisingly, only 7 of the 11 validated NPC components in *Plasmodium berghei* share significant primary sequence homology with any protein in *Toxoplasma.* In fact, we were unable to find potential orthologs to most *Toxoplasma* nucleoporins in *Plasmodium* (Data S2), highlighting the level of divergence even within phylum Apicomplexa. Even within alveolates and other eukaryotes we find orthologs of only a core set of *Toxoplasma* Nups and the majority remain oblivious to sequence based orthology. Much of this apparent lack of orthology is likely due to the detection limits of sequence-based methods. However, the structural differences between the *Toxoplasma* NPC and that of other eukaryotes [5,7] and the lack of clear orthologs to proteins that are well-conserved outside of Apicomplexa suggest the potential for a large amount of specialization.

One of the major sources of difference we identified were in both the cytoplasmic and nucleoplasmic components for mRNA export. While the core scaffolds for both complexes appear well conserved, different Alveolate lineages are missing specific components of each. For instance, while Rae1/Gle2 is conserved in Apicomplexa and dinoflagellates it is missing in ciliates. However, the core of the cytoplasmic mRNA complex appears well conserved, and we confirmed Nup139b/TG*_297830 and Gle1/TG*_259172 as essential components that to this structure by UExM.

The TREX-2 complex acts in the nucleoplasm to shuttle mRNA to the NPC for export [71–74]. We identified and functionally validated the core component of this complex as Sac3/TG*_211700. In yeast, TREX-2 is tethered to the NPC by ScNup1 [71] and ScCdc31/Centrin-3 [50]. *Toxoplasma* has a clear Centrin-3 ortholog [51], and we identified Nup145/TG*_203780 as a potential ortholog of Nup1. However, unlike in yeast, loss of the *Toxoplasma* proteins had no effect on mRNA export (Figure 5). This can be partially explained by the fact that ScNup1 serves in yeast to anchor TREX-2 to the nuclear basket, a structure that is largely missing in *Toxoplasma* [7]. Indeed, a recent study demonstrated that the human TREX-2 complex makes intimate contacts with both the Y-shaped complex and nuclear spokes [74], which are composed of Mlp1/TPR. While conserved in other alveolates, Mlp1 and other nuclear basket proteins are missing in Apicompexa (Data S4). This suggests that the TREX-2 complex must be anchored through a different mechanism than in opisthokonts. These findings are in stark contrast to a recent analysis of the NPC in another divergent parasite from a different phylum, *Trypanosoma brucei [60]*. Clear orthologs to many yeast nucleoporins, including Mlp1/Mlp2, have been identified in trypanosomes [15,25,60]. Similarly, while several *Trypanosoma-*specific components have been identified in the mRNA export machinery, both the cytoplasmic-facing and nuclear machinery appears well conserved in trypanosomes, including each of the components of the TREX-2 complex [25]. Notably, our bioinformatic analysis identified potential orthologs of both the nuclear basket protein Mlp1 and the TREX-2 component Sus1 in other Alveolate lineages, including Chromerids (Data S2-4). It therefore seems that any apicomplexan specialization of these complexes is unusual to the phylum.

By examining markers for subcomplexes on opposite sides of the nuclear envelope, we identified *Toxoplasma* proteins required for the stability of specific structures within the NPC (Figures 8-9, 12). We found that degradation of Nup281/TG*_226950 resulted in displacement of Nup139b/TG*_297830 without affecting Nup302C. Intriguingly, Nup281/TG*_226950 knockdown did not immediately affect mRNA export, suggesting it is not a directly involved in this process (Figure 7B, S11D). Our UExM data show Nup281/TG*_226950 localizes near Nup139b/TG*_297830 on the cytoplasmic face of the NPC, and therefore suggest Nup281/TG*_226950 is instead a parasite-specific scaffold that anchors the cytoplasmic platform to the NPC. We also found that degradation of Nup145/TG*_203780 resulted in the detachment of Nup302C from the NPC, suggesting that Nup145/TG*_203780 is a critical component of the Y-shaped complex, and perhaps an ortholog of ScNup1. Because ScNup1 is also part of the nuclear basket in yeast, Nup145/TG*_203780 may provide a useful hook to examine what residual structure of the nuclear basket is preserved in *Toxoplasma*.

We found that the likely adapter protein, Nup433/TG*_214600, is required for efficient assembly of NPCs. Similarly, loss of function of the yeast adapter proteins ScNup157 and ScNup170 [57] results in the gradual depletion of functional NPCs with each cellular division. Thus our data suggest that the underlying paradigm of NPC assembly is conserved in *Toxoplasma* in spite of its divergent architecture. Note that ScNup157/170 are paralogs. Deletion of either gene does not affect yeast growth, though the double mutant is synthetic lethal [57]. Based on homology and secondary structure prediction (Figure 2, Data S2), we predict that Nup327/TG*_258970 is an ortholog of these proteins and found that it is non-essential. It is therefore possible that a second protein in *Toxoplasma* assists Nup327/TG*_258970 in its function. Such overlapping essential function is a common theme in a massive structure such as the NPC [41,57], which helps explain the resistance of the complex to loss of individual components.

The subcomplexes of the NPC were originally characterized biochemically – by identifying proteins that co-immunoprecipitated in different conditions. The tremendous strides in our understanding of the NPC three-dimensional structure have required an integrative approach, combining high-resolution structures of individual subcomplexes, crosslinking-mass spectrometry, and cryo-electron microscopy and tomography of intact NPCs [6,7,17,75–79]. All of these experiments, as with the reverse genetics that has enabled functional characterization of the NPC’s diverse functions, have required knowledge of the proteins involved. As we have now validated a large number of *Toxoplasma* Nups, we expect our data to empower future studies of the peculiarities of the structure and cellular functions of the NPC in this divergent organism.

## Materials and Methods

### PCR and Plasmids

The primers used in this study for generation of plasmids are listed in Data S5. All PCRs were done using Phusion polymerase (NEB; New England Biolabs) and plasmids were assembled using Gibson master mix (NEB).

### Parasite Culture and transfections

*Toxoplasma* tachyzoites were maintained on confluent HFF (Human foreskin fibroblasts) monolayers. The HFFs were cultured in DMEM (Dulbecco’s modified Eagle’s medium) supplemented with 10% FBS (fetal bovine serum) and 2 mM glutamine. The Nup302^BioID2-3xHA^, Nup134^BioID2-3xHA^ and Nup68^BioID2-3xHA^ parasites were generated by transfecting ∼30 μg of linearized plasmid containing ∼1500 bp of targeting sequence in frame with the BioID2 and 3xHA tags in the *RHΔku80Δhxgprt* strain [80]. Selection was performed using 25 μg/mL mycophenolic acid and 50 μg/mL xanthine (MPA/Xanthine) in the culture medium. Clonal parasite lines were obtained by infecting a 96 well plate with 1 parasite per well and screening by immunofluorescence (IFA) for HA signal using anti-HA antibody (3F10, Roche/Sigma). For generating parasites with tagged NPC markers, linearized plasmid containing homologous regions of either Nup302, TG*_297830, or TG*_211700 in frame with 6xTy1 tag, or TG*_258850 in frame with a 3xFLAG tag were transfected into the *RHΔku80Δhxgprt* parasites expressing OsTir1 driven by the *gra1* promoter. Selection was performed with Zeocin (Bleomycin). Briefly, parasites from a highly infected T25 were released using a syringe and a 27 gauge needle, filtered and centrifuged at 300xg to collect parasites. The parasites were resuspended in 500 μL of HBSS (Hank’s balanced salt solution) containing 25 μM Zeocin. The parasites were incubated at 37°C for 1 hour and passed onto a fresh T25 of HFFs for outgrowth before cloning by limited dilution, as above. The AID-3xHA-tagged strains for all selected candidates were generated by transfecting the *RHΔku80Δhxgprt* Ostir1/Nup302C^6xTy1^ strain with linearized plasmids containing ∼700 to ∼1500 bp of targeting sequence in frame with the 3xHA and AID tags. These strains were used for initial verification of candidates and determining essentiality of the candidate proteins. Essential Nups were also tagged in the background of OsTIR1/TG*_297830^6xTy1^. The OsTIR1/TG*_211700^6xTy1^ strain was transfected with linearized plasmid to tag TG*_297830 with 3xHA-AID tag. Similarly, to test nucleocytoplasmic transport, the OsTIR1/TG*_258850^3xFLAG^ parasites were tranfected to tag TG*_214600, TG*_275490, and TG*_313430 with 3xHA-AID tag.

### Proximity Biotinylation and mass spectrometry

Nup302-BioID2-3xHA, Nup134-BioID2-3xHA, and Nup68-BioID2-3xHA parasites were passed into 2× 15cm dishes each and grown in presence of 150 μM biotin (Sigma-Aldrich) for 40 h. The infected HFF monolayer was washed with Phosphate Buffered Saline (PBS) 3× to remove any extracellular biotin. The parasites were released from the host cell using a syringe and a 27 gauge needle and harvested by centrifugation. The parasite pellet was resuspended and washed 3× with 50 mL PBS. The parasite pellet was then resuspended in 2.5 mL of 1x RIPA buffer and incubated at 4°C for 2 hours to ensure lysis. In order to remove remaining free biotin, each sample was buffer exchanged 2× using a pre-equilibrated PD-10 Desalting column (GE Healthcare). The biotinylated proteins were separated from the lysate using magnetic streptavidin resin (NEB). The resin was washed 2× with RIPA buffer, 1× with 20 mM Tris-HCl pH 7.5, 1 % SDS, 1× with each: 10 mM Tris-HCl pH 8.1, 250 mM LiCl, 0.5% NP40, 0.5% Sodium deoxycholate, 1 mM EDTA; 50 mM Tris-HCl pH 7.4, 50 mM NaCl, and finally 2× with PBS. Biotinylated proteins were eluted with 60 mM Tris-HCL pH 6.8, 4% SDS, 16.6% Glycerol and 10 mM β-Mercaptoethanol (2x SDS loading dye). Elutions were loaded on precast Mini-PROTEAN TGX Stain-Free Precast Gels (BioRad) and run at 200V for 6 mins. The gel was stained with gel code blue (invitrogen) and stained bands were cut and processed for mass spectrometry analysis.

For mass spectrometry analysis, individual samples were digested overnight with trypsin (Pierce) following reduction and alkylation with DTT and iodoacetamide (Sigma–Aldrich). The samples then underwent solid-phase extraction cleanup with an Oasis HLB plate (Waters) and the resulting samples were injected onto an Orbitrap Fusion Lumos mass spectrometer coupled to an Ultimate 3000 RSLC-Nano liquid chromatography system. Samples were injected onto a 75 μm i.d., 50-cm long EasySpray column (Thermo) and eluted with a gradient from 0-28% buffer B over 60 min at 250 nL/min. Buffer A contained 2% (v/v) ACN and 0.1% formic acid in water, and buffer B contained 80% (v/v) ACN, 10% (v/v) trifluoroethanol, and 0.1% formic acid in water. The mass spectrometer operated in positive ion mode with a source voltage of 2.2 kV and an ion transfer tube temperature of 275 °C. MS scans were acquired at 120,000 resolution in the Orbitrap and up to 10 MS/MS spectra were obtained in the ion trap for each full spectrum acquired using higher-energy collisional dissociation (HCD) for ions with charges 2-7. Dynamic exclusion was set for 25 s after an ion was selected for fragmentation.

Raw MS data files were analyzed using Proteome Discoverer v2.2 (ThermoFisher Scientific), with peptide identification performed using Sequest HT searching against the *Toxoplasma* GT1 and ME49 proteomes from ToxoDBv68 [62]. Fragment and precursor tolerances of 10 ppm and 0.6 Da were specified, and three missed cleavages were allowed. Carbamidomethylation of Cys was set as a fixed modification, with oxidation of Met, methylation of Lys and Arg, demethylation of Lys and Arg, trimethylation of Lys, acetylation of Lys, and phosphorylation of Ser, Thr, and Tyr set as a variable modification. The false-discovery rate (FDR) cutoff was 1% for all peptides.

### Immunofluorescence and Microscopy

HFFs were plated on 24 well plates containing 1.2 cm (diameter) glass coverslips and grown to confluency. The confluent coverslips were infected with parasites and incubated overnight or to a fixed time-point. The infected coverslips were washed with PBS and fixed with 4% paraformaldehyde + 4% sucrose in PBS at room temperature for 15 min. The coverslips were washed 3× with PBS before adding 0.1% Triton X-100 in PBS (PBST) for 20 min to permeabilize the cells. The coverslips were washed 3× with PBS and blocked for 45 min with 3% Bovine Serum Albumin (BSA) in PBS in a humidity chamber. The coverslips were then incubated with primary antibody overnight at 4°C and washed 3× with 0.1% Triton X-100 in PBS with 5 min incubation between each wash. The coverslips were then treated with secondary antibody conjugated with Alexa-flour (Molecular probes) for 40 min and washed 3×, with 5 min incubation between each wash. The coverslips were then treated with Hoechst for 15 min where needed, washed with PBS and then mounted on a glass slide using mounting medium (Vector Laboratories). The cells were imaged on either a Nikon A1 laser scanning confocal microscope with a 60× oil immersion 1.42 NA objective or with a Nikon Ti2E wide-field microscope with a 100× oil immersion 1.45 NA objective using Nikon Elements. Primary antibodies used in this study are rat anti-HA (Sigma-Aldrich; 1:1,000 dilution), rabbit anti-Tg-β-tubulin (1:10,000 dilution), mouse anti-Ty1 (Sigma-Aldrich; 1:10,000). The secondary antibodies used in the study are Goat anti-Rabbit IgG, Alexa Fluor™ 647 (Thermo; # A-21245), Goat anti-Mouse IgG, Alexa Fluor™ 555 (Thermo, # A-21424) and Goat anti-Rat IgG, Alexa Fluor™ 488 (Thermo, #A21208).

### Ultra Expansion Microscopy (U-ExM)

UExM protocol was adapter from Liffner and Absalon [43,81]. Coverslips were prepared and fixed as mentioned in the immunofluorescence and microscopy section. Crosslinking solution was prepared by mixing 40 μL of monomer solution (19% Sodium Acrylate, 10% Acrylamide, 0.1% Bisacrylamide, in 1x PBS) with 5 μL TEMED and 5 μL APS. 35 μL of crosslinking solution was placed immediately on a parafilm sheet kept in a humidity chamber that was placed on ice. The fixed coverslip was placed inverted on top of the crosslinking solution and kept on ice for 5 mins. The coverslips were then incubated at 37°C for 30 min for gelation. The gels were removed from coverslips in a 6 well plate with 2mL of denaturation buffer (50 mM Tris HCl pH 9.0, 350 mM sodium dodecyl sulphate (SDS), 200 mM NaCl) with rotation at room temperature for 15 mins. The gels were separated and transferred to a 2 mL tube, filled with denaturation buffer, and incubated at 95°C for 90 mins. The gel was allowed to cool and then expanded in a 10 cm dish by washing with 25 mL of MilliQ water for 30 min at room temperature, thrice. The gel was then washed with PBS twice for 15 min each time. Gels were moved to a clean 6 well plate and blocked with 3% BSA in PBS for 30 minutes and then incubated overnight with primary antibody in 3% BSA. The next day, the gel was washed 3× for 10 min with 1× PBST and then incubated with a secondary antibody in 3% BSA for 150 min. The gel was washed 3× for 10 min with 1× PBST. The gel was expanded by washing thrice with MilliQ water for 30 min each in a 10 cm dish. In the last wash, Hoechst was added to stain the nuclei. The gel was then transferred to a new 10 cm dish and the diameter measured to calculate the expansion factor. The expanded gel was cut and transferred to a glass-bottom dish for imaging. All expanded gels were imaged on Zeiss LSM900 equipped with Airyscan array detector in super resolution mode. Images were taken with DAPI (Ex:355 Em:465), Alexa Fluor 488 (Ex:493 Em:517), and Alexa Fluor 555 (Ex:553 Em:568) laser lines using a Plan-Apochromat 63x/1.40 Oil DIC M27 objective. The pixel size was 35.3 nm and the step size was 0.14 μm. Acquired raw images were deconvoluted with Zeiss in-built Zen deconvolution tool. The images were further analyzed with Fiji/ImageJ [82].

For counting the TG*_297830 puncta, we used 3D object counter in Fiji/ImageJ with a minimum object size of 75 voxels. The puncta were reviewed and audited manually and aggregates were removed. The cropped image was changed to an 8-bit image and smoothened with a gaussian blur (sigma = 2.0). We applied a binary mask on the smoothed image and proceeded with the fill holes function to connect the nuclear volume. We then used the 3D object counter to calculate the volume and surface area of the nucleus. For preparing representative images, the drift between stacks was corrected using MultistackReg [83] and then the maximum intensity projections were generated in Fiji/ImageJ [82].

### Plaque Assays

To measure the survival of parasites upon knockdown of candidate Nups, 200 or 100 parasites from freshly syringe released T25 of a respective clonal population were added to a monolayer of HFFs in a 6-well plate in IAA or vehicle (500 μg/mL; dissolved in 100% ethanol). After 8 or 10 days, the HFF monolayer was fixed with methanol and stained with crystal violet. All plaque assays were performed in n=3 biological replicates with n=3 technical replicate per biological replicate. For the candidate Nups that showed plaques upon knockdown, the plaque areas were measured using Fiji/ImageJ [82]. Significance was determined by unpaired two-tailed Student’s t-test.

### Parasite Replication and parasitophorous vacuole (PV) size measurement

Parasite replication was quantified by counting the number of parasites per vacuole. Parasites with AID-tagged Nups were passed on coverslips and allowed to infect for 2 hours. The media was changed to ±IAA after 2 hours of infection and the coverslips were fixed 16 hours after media change. IFA was performed as above and the parasites were visualized with anti-HA, anti-Tub, anti-Ty1 antibodies, and Hoechst stain. The resulting images were analyzed using Fiji/ImageJ [82]. For each replicate, 100 vacuoles per replicate were analyzed to count the number of vacuoles containing 1, 2, 4, or 8+ parasites in ±IAA conditions for a total of n=3 biological replicates. Significance between the ±IAA conditions was determined by Kolmogorov–Smirnov test.

The number of inviable parasites was measured based on the tubulin staining for TG*_313430^AID^, TG*_203780^AID^, TG*_275490^AID^, and TG*_214600^AID^ at 24 h. The vacuole size was measured for TG*_214600^AID^ at 24h and 36h. For both measurements, the coverslips with confluent HFF monolayers were infected with the clonal parasite line with a MOI of 0.1 and allowed to infect for 2 hours before changing the media to ±IAA. The coverslips were fixed at the timepoints mentioned for the respective experiments. The parasites were stained with anti-Gra1, anti-tubulin, and Hoechst stain. Maximum intensity projections of widefield images were used for measuring the vacuole size and for assessing the inviable parasites. The tubulin signal was manually assessed as normal or abnormal for n=30 parasites per biological replicate. Data from three biological replicates were plotted as a normalized stacked bar graph and significance between the ±IAA conditions was calculated by Kolmogorov–Smirnov test. For measuring the vacuole size, the largest dimension of GRA1 signal was measured in Fiji/ImageJ [82] for each time point. Data from three biological replicates, each an average of n>20 vacuoles, were plotted on a graph against their respective time points. Significance was calculated using unpaired two-tailed Student’s t-test at each time point.

### mRNA export assays

We adapted a published fluorescence *in-situ* hybridization protocol [84]. To measure the defect in mRNA transport, parasites from a clonal population of AID tagged candidate parasites were allowed to infect a coverslip with a confluent HFF monolayer. The parasites were allowed to grow for 16 hours before changing the media to ±IAA for 2 hours. The HFF monolayer was fixed and permeabilized as mentioned in the Methods section (Immunofluorescence and Microscopy). Following permeabilization, the cells were then incubated in a 1:1 mixture of PBS + 0.3% Triton-X and Pre-Hybe buffer (50% De-ionized Formamide [Roche], 5x SSC [saline-sodium citrate], 1mg/ml yeast RNA [Sigma], 1% Tween-20 [Sigma, from 10% stock]) for 10mins at RT. After transferring the coverslips to a humidity chamber, the cells were incubated in Pre-Hybe buffer for 30 min at 37°C. Post incubation, 200 ng of fluorescein tagged oligo(dT) (60-mer) mixed in Hybe buffer (50% De-ionized Formamide [Roche], 10% Dextran Sulfate [Sigma], 5x SSC, 1mg/ml yeast RNA [Sigma], 1% Tween-20 [Sigma, from 10% stock]) was added to each coverslip and further incubated overnight at 37°C. Cells were then washed with pre-warmed Wash Hybe buffer (25% Standard Formamide [Roche], 3.5x SSC, 0.5% Tween-20 [Sigma], 0.05% Triton X100 [Sigma]) 2× for 15 min each at 37°C. The cells were further washed an additional 10× with pre-warmed 1xPBS for 10 min each at 37°C. The cells were further stained with Hoechst for 15 min at RT before mounting and imaging. The cells were imaged on a Nikon A1 laser scanning confocal microscope with a 60× oil immersion 1.42 NA objective by using Nikon Elements software. A single slice of a confocal image from the center (in z) of the nucleus was used to quantify the poly(dT) signal. Hoechst signal was used to identify nuclei. Nuclear mean fluorescein intensity was quantified from background subtracted images. N=6 biological replicates were quantified, each with a minimum n=20 technical replicates. Significance was calculated by unpaired two-tailed Student’s t-test.

### NPC marker mislocalization assay

We assessed NPC marker mislocalization for all the essential Nups with both TgNup302C^6xTy1^ and TG*_297830^6xTy1^ NPC markers. Clonal parasites were passed onto a coverslips with a confluent HFF monolayer and the media was changed to ±IAA after 2 hours to allow for invasion. Coverslips were fixed after 8 hours of ± IAA treatment. The parasites were visualized with anti-HA, anti-Tub, anti-Ty1 antibodies, and Hoechst stain. For making the scoring rubric of phenotypes and assessing the mislocalization of both NPC markers, we used maximum intensity projections of widefield images having the Ty1 and tubulin signal. 3 images per essential Nup per NPC marker per condition were used for making the rubric. The remaining images were blinded and 7 images per strain per NPC marker per condition were assessed for NPC marker localization defects for n=5 biological replicates. For determining NPC marker mislocalization (TgNup302C^6xTy1^, TG*_297830^6xTy1^, and TG*_211700^6xTy1^) in the candidates that showed mRNA accumulation (TG*_211700 and TG*_297830) upon their knockdown, we incubated the parasites for 2 hours in normal media followed by a 2 hour incubation in ±IAA media. The same strategy was used to access the NPC marker mislocalization at 2 hours for an n=3 biological replicates. Similarly, we analyzed TG*_203780^AID^/Nup302C^6xTY1^ and TG*_226950^AID^/ TG*_297830^6xTy1^ parasites with a 2 hour infection and IAA treatment for 2, 4, and 6 hours with n=3 biological replicates per condition. Significance was calculated by Welch’s t-test and the p-values were adjusted for multiple tests using Benjamini–Hochberg false discovery rate correction.

### Nucleocytoplasmic transport Assays

To test the defect in nucleocytoplasmic transport, HFF monolayers on coverslips were infected with AID tagged parasites expressing the NCT marker TG*_258850^3xFLAG^ for 2 h and then treated with IAA for 8 h. The coverslips were fixed and IFA stained with Hoechst, rat anti-HA, mouse anti-FLAG and rabbit anti-Tubulin. The coverslips were imaged on the Nikon A1 laser scanning confocal microscope with a 60× oil immersion 1.42 NA objective by using Nikon Elements software. The Z-stacks, with four channels (405nm, 488 nm, 561 nm and 640 nm), were taken with a step size of 0.5 µm, pinhole of 1 AU at TRITC (561 nm) and a resolution of 19.3087 pixels per micron (1024×1024). These settings were maintained across all acquisitions for all images. The stacks were opened in Fiji/ImageJ [82] and the background was subtracted with a rolling ball radius of 40 pixels. Hoechst signal was used to mark the nucleus and tubulin signal was used to mark the parasite. The FLAG signal was quantified for both the nucleus and the parasite, and reported as a ratio of nuclear over total average intensity. This values were used for comparing the ± auxin conditions graphically and for calculation of average values to finally compare them using an unpaired Student’s t-test.

### Bioinformatic and structural analysis

*Toxoplasma gondii* (GT1) and *Plasmodium berghei* (ANKA) Nup sequences were obtained from VEuPathDB [62] and the yeast sequences were obtained from UniProt [85]. BLAST [66] and PANTHER [67] searches were performed on the NCBI webserver with an e-value cutoff of 10. Reciprocal BLAST searches were performed for all hits of *Toxoplasma* and *Plasmodium* searches below a cutoff of 1e-2. For the Foldseek [68] searches, Alphafold models of *Toxoplasma*, *Plasmodium* and yeast were downloaded from the Alphafold database [86]. If the models were not available in the database, Nup sequences were used to construct models using AlphaFold3 webserver [68]. These models were used to search for orthologs on the Foldseek webserver. The secondary structure prediction was performed using s4pred [88]. The coiled-coil prediction was performed using Deepcoil2 [89]. Transmembrane domain prediction was performed using DeepTMHMM [90].

We used ChimeraX [91] to visualize, analyze and generate images showing the cryo-ET maps from *Toxoplasma* (EMD-44381), Yeast (EMD-44377), and the integrative models (PDB: 9A8M, 9A8N, 9A0F) [6,7]. We generated the TREX-2 complex model using AlphaFold 3 webserver [87]. The stoichiometries for the TREX-2 complex components were based on PDB: 4MBE (yeast Sac3:Sus1:Cdc31:Nup1 complex) [45], 3FWC (yeast Sac3:Sus1:Cdc31 complex) [72] and 8U8D (yeast Sac3:Thp1:Sem1 complex) [46]. ChimeraX was used to visualize the model and generate image for the figure.

To predict the subcomplex membership in Figure 6, we combined the above bioinformatic data with the following rubric: (i) membrane Nups are defined by one or more transmembrane domains. (ii) Central channel Nups have an FG repeat region and a helical coiled-coil region. (iii) The adapter Nups and the Y-shaped complex include proteins with an α-solenoid fold. In addition, the adapter Nups, the Y-shaped complex, and the cytoplasmic mRNA export platform include Nups containing a β-propeller followed by a helical solenoid (in the adapter and Y-shaped complex) or coiled-coil domains (mRNA export platform). (iv) Linker Nups are predicted to be intrinsically disordered and usually contain FG repeats. (v) Most components of the nuclear basket are intrinsically disordered, other than Mlp1/2 which are coiled-coil proteins.

To identify potential orthologs of well-folded human and yeast Nups we used HMMERv3.4 to search the core SAR genomes from EukProtV3 [63]. We included select genomes from Amoebazoa, animals, fungi, trypanosomes, and plants as outgroups. The top hit for each organism was added to a list of unaligned sequences which were then aligned with ClustalO [92]. Phylogenetic trees were estimated by IQTREE2 using SH-aLRT [93] and ultra-fast bootstrapping [94] and the model selected automatically (support values are reported as “SH-aLRT/bootstrap” in the resulting trees). Accession numbers are in Data S3 and alignments and phylogenetic trees are in Data S4.

### Figure generation

All graphs were plotted using the matplotlib and seaborn modules in Python. Statistics were calculated using scipy.stats. All microscopy images were generated using Fiji/ImageJ [82]. All images were annotated and organized using Inkscape v1.4 to generate final figures.

**Table 1:**
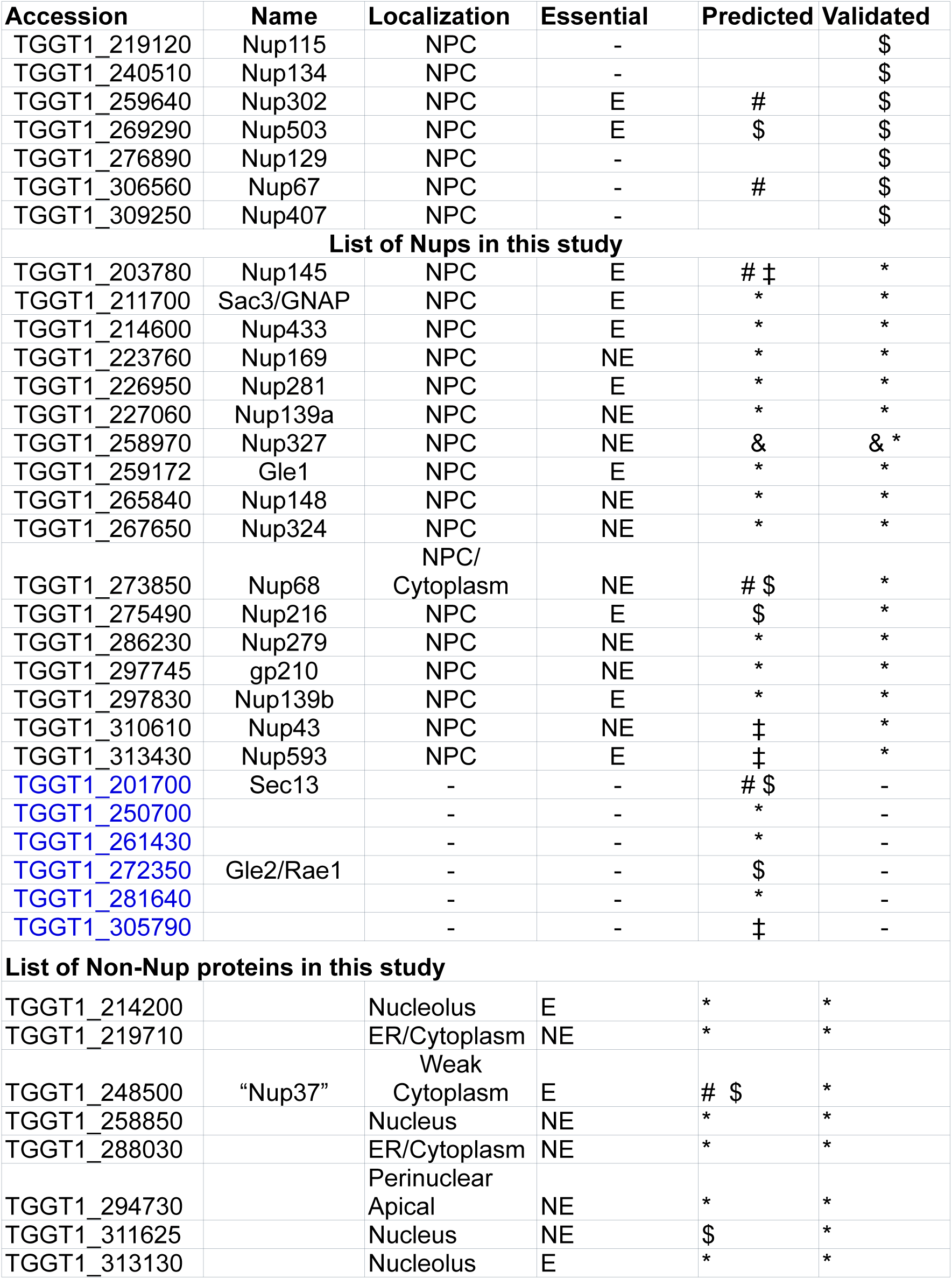
Overview of predicted *Toxoplasma* NPC components validated in this study or elsewhere. Nup names are based on orthology (when non-Nup designated) or predicted molecular weight (per the standard Nup nomenclature). Experimentally validated localizations are indicated as well as the study that demonstrated the localization (* This study; # Neumann [27]; $ Courjol [20]; ‡ Kehrer [18]; & Ovciarikova [21]; + Frankel [95]).

**Table 1:**
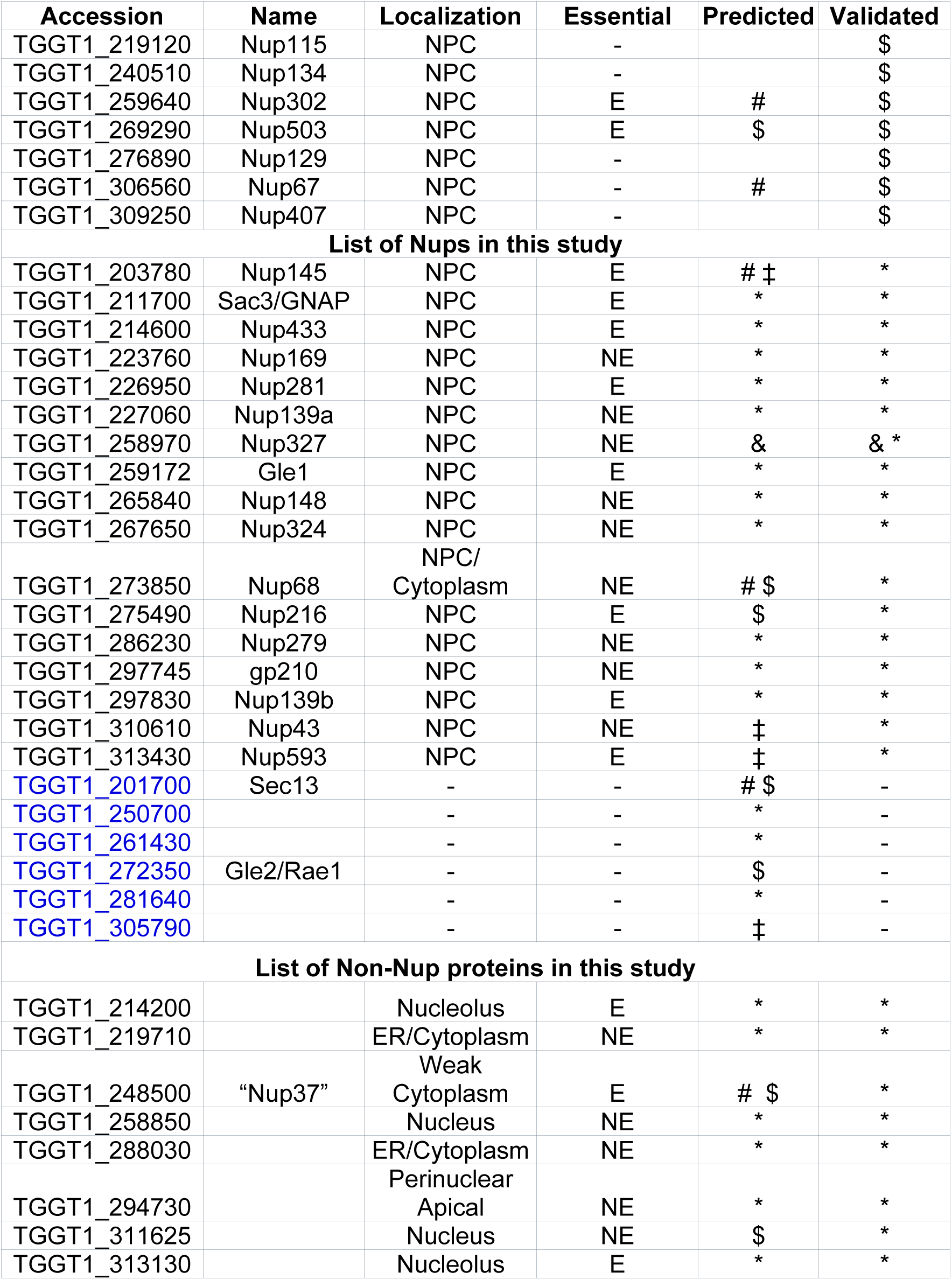
Overview of predicted *Toxoplasma* NPC components validated in this study or elsewhere. Nup names are based on orthology (when non-Nup designated) or predicted molecular weight (per the standard Nup nomenclature). Experimentally validated localizations are indicated as well as the study that demonstrated the localization (* This study; # Neumann [1]; $ Courjol [2]; ‡ Kehrer [3]; & Ovciarikova [4]; + Frankel [5]).

## Data Availability

Mass spectrometry data are available from the MassIVE database with accession MSV000098791.

## Supporting information

SI Data S1 BioID

SI Data S2 Bioinformatic analysis

SI Data S3 accessions used

SI Data S4 - alignments and phylogenies

SI Data S5 - List of primers

## Acknowledgments

We acknowledge the Proteomics Core Facility at University of Texas Southwestern Medical Center for mass spectrometry analysis. Molecular graphics and analyses performed with UCSF ChimeraX, developed by the Resource for Biocomputing, Visualization, and Informatics at the University of California, San Francisco, with support from National Institutes of Health R01-GM129325 and the Office of Cyber Infrastructure and Computational Biology, National Institute of Allergy and Infectious Diseases. We thank the Collins lab at UTSW for their help with the FISH protocol and allowing us to use their Zeiss Airyscan microscope. We thank Sabrina Absalon for help with the UExM. We thank Ben Weaver and Vasant Muralidharan and members of the Reese lab for helpful comments on the manuscript. M.L.R. acknowledges funding from the Welch Foundation (I-2075-20240404) and a Burroughs Wellcome Foundation Investigators in Pathogenesis of Infectious Disease award (G-1021959).

---

**Figure S1:**
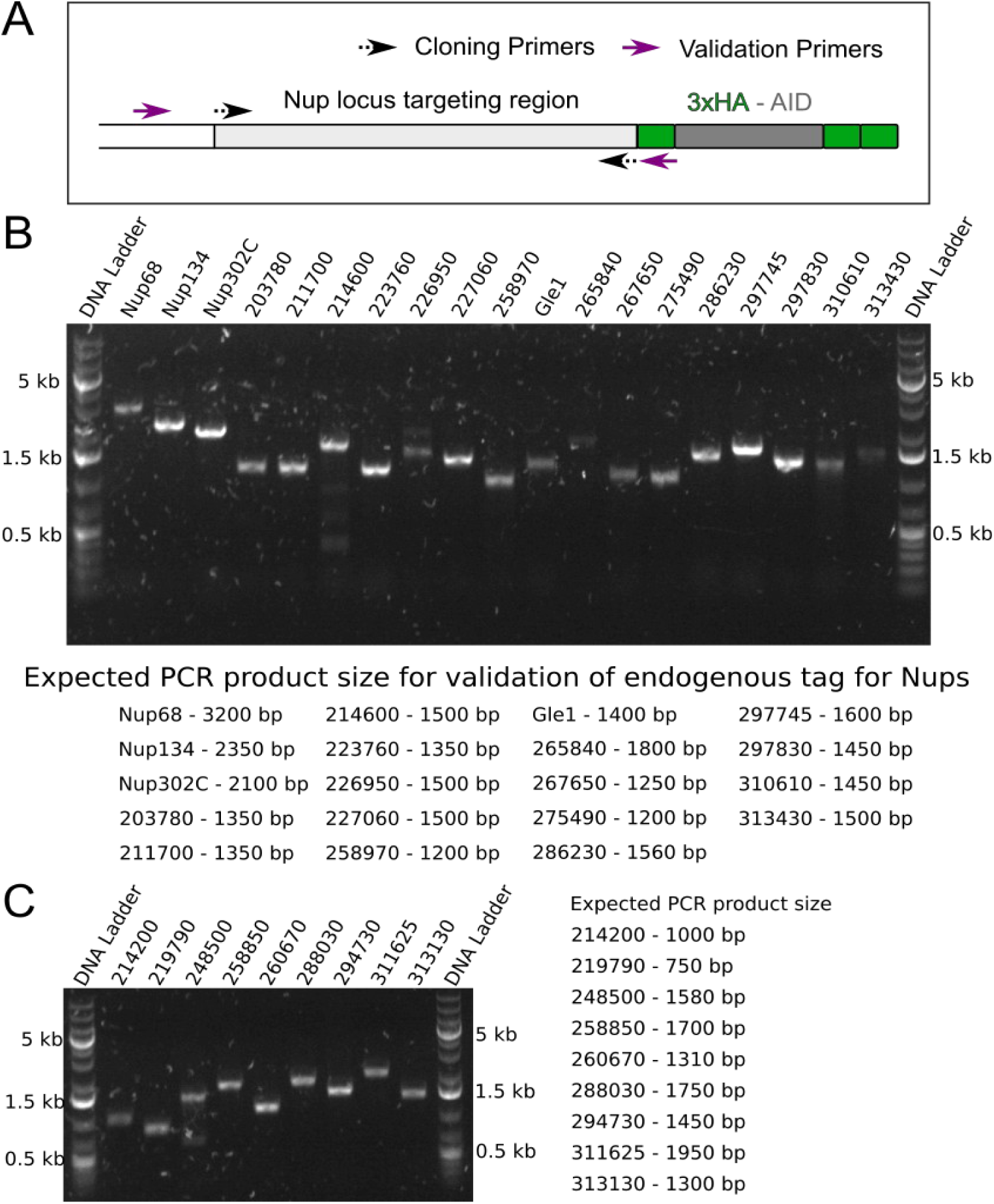
Endogenous tag for candidate proteins was verified by PCR. (A) Cartoon representation of the tag verification strategy highlighting that the validation primers bind outside the targeting region. PCR amplicons using the gDNA from respective strains for mentioned Nups (B) and non-Nups (C) were resolved on an agarose gel to confirm the endogenous 3xHA-AID tag. The expected sizes are mentioned below the gel image for Nups, and adjacent to the gel image. GeneRuler 1 kb Plus DNA Ladder is used.

**Figure S2:**
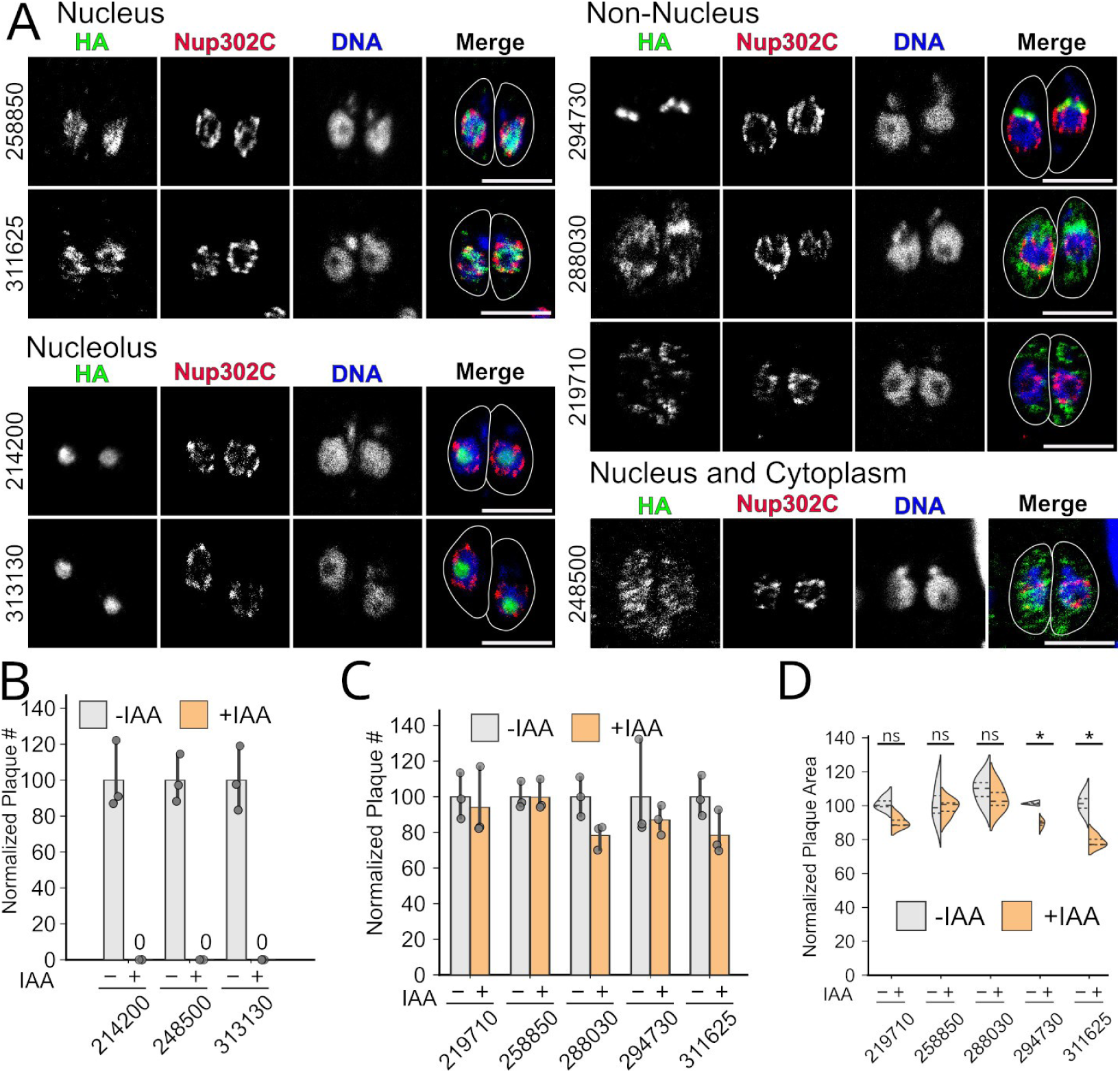
A subset of candidates did not localize to the NPC. (A) Intracellular parasites in which the indicated candidates (for clarity, only the numerical identifiers are indicated, i.e., TG*_######) had been tagged with AID-3xHA were stained with antibodies recognizing HA (green), Ty1 (red; Nup302C), and Toxoplasma β-tubulin (not shown; used to create outline). DNA was stained with Hoechst (blue). Images are 1 airy-unit confocal slices (∼0.75 μm). All scale bars are 5 μm. Note lack of localization to NPC (red) for all candidates. While some of these non-Nup candidates were unable to form plaques when grown in +IAA (B), others showed no loss in plaque efficiency (C). (D) A subset of these non-essential proteins showed mild fitness cost upon growth in +IAA, per a reduction in plaque area. ns, not significant; *, p<0.05; n=3 biological replicates two-tailed unpaired Student’s t-test.

**Figure S3:**
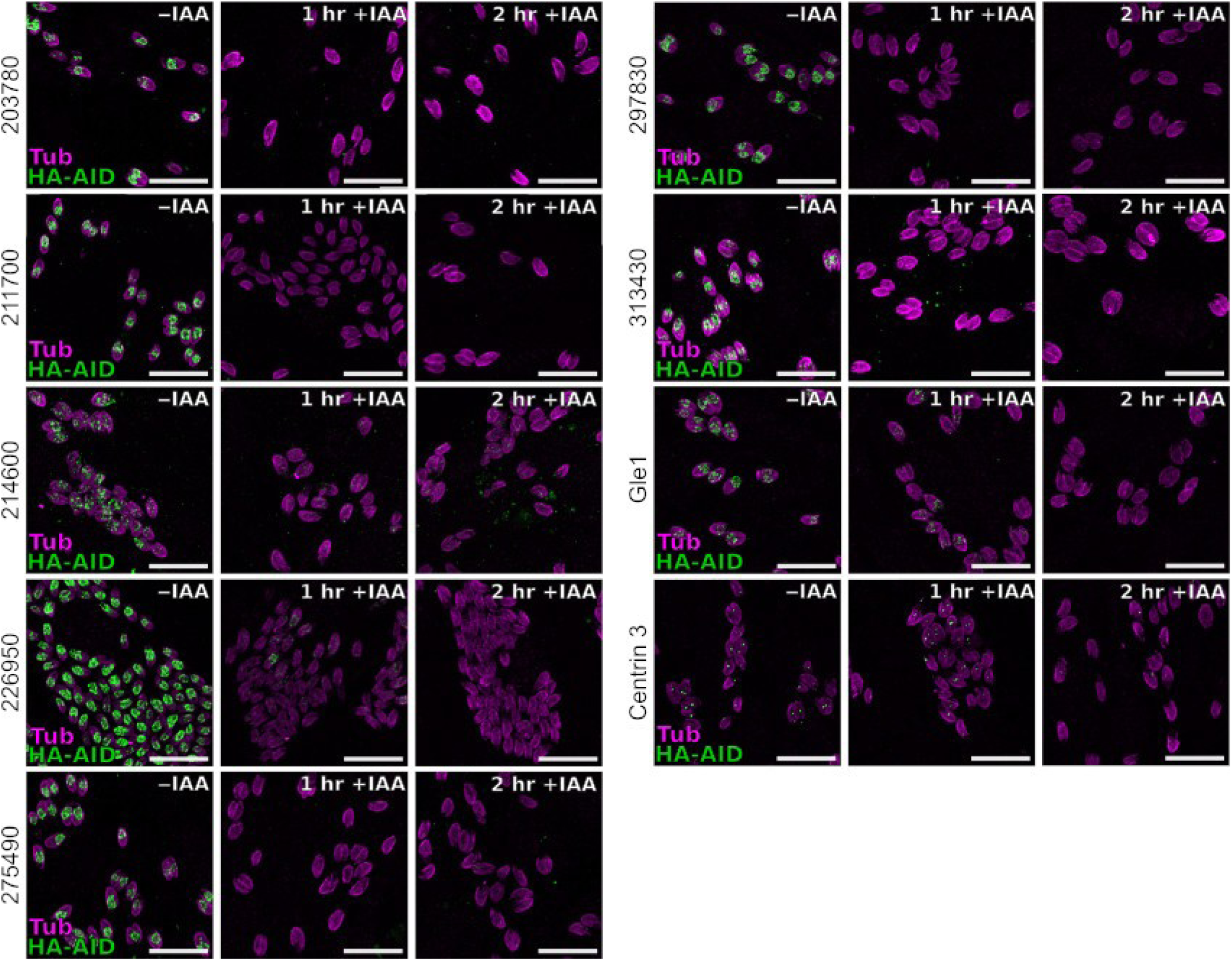
Essential Nups are knocked down by 2 hours of IAA treatment. Maximum intensity projection of confocal stacks of parasites grown in ±IAA media for 1 or 2 hours show complete knockdown of the respective Nup at 2 hours of IAA treatment. Scale bars are 20 µm.

**Figure S4:**
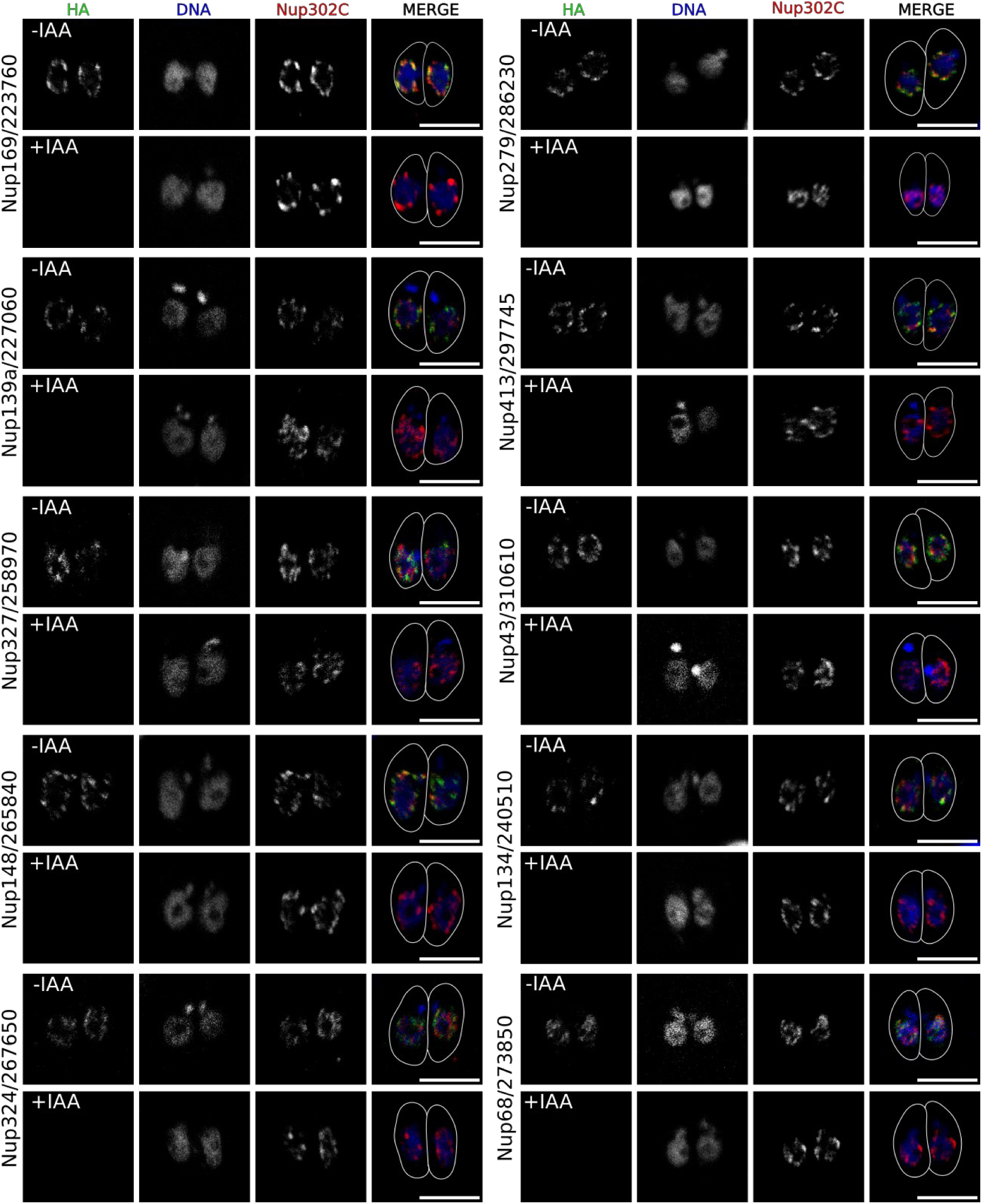
Knockdown confirmation for Non-Essential Nups. Single slices of confocal micrograph of parasites treated with ±IAA for 6 hours. Parasites in +IAA have no remaining AID-3xHA signal. Scale bars are 5 µm.

**Figure S5:**
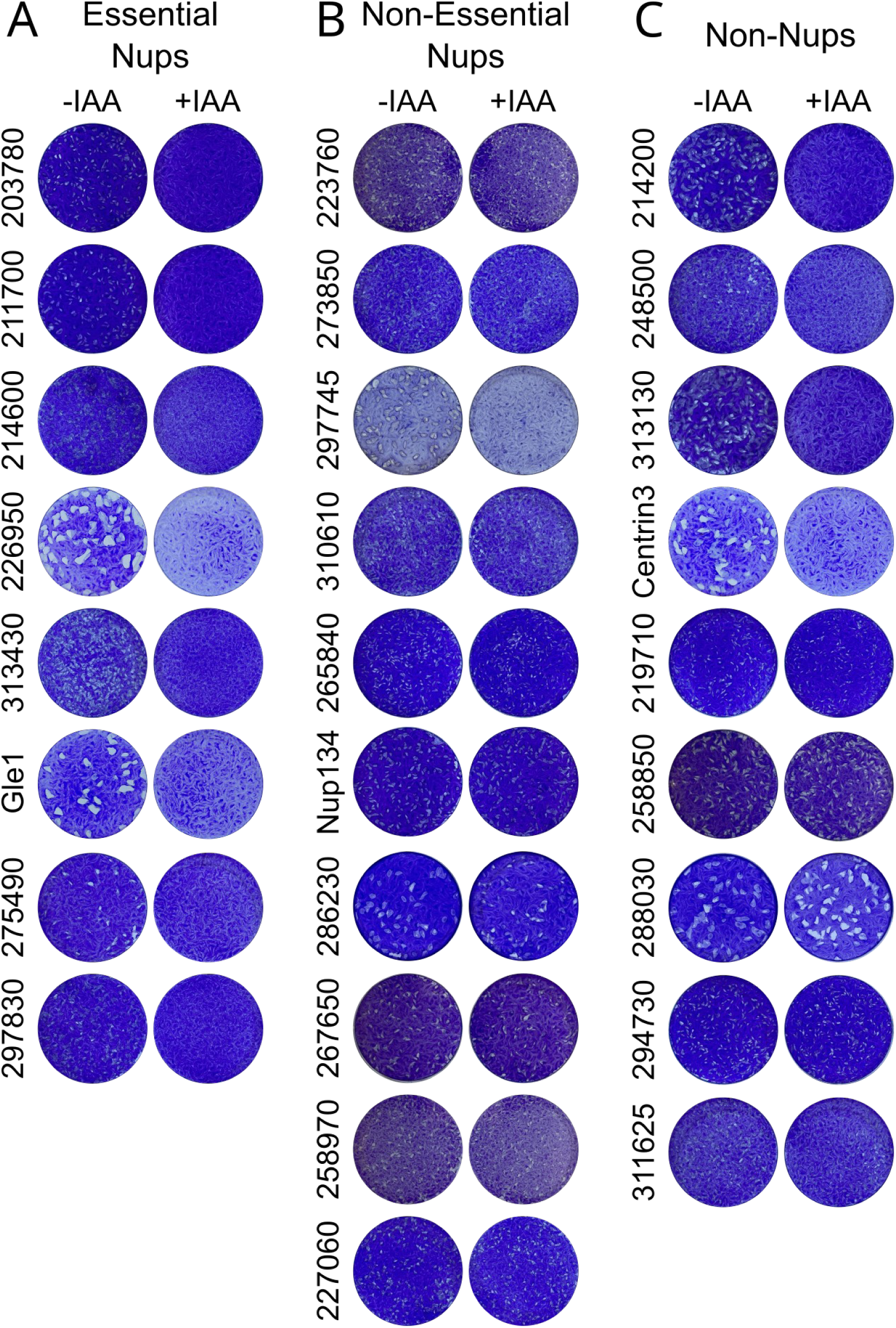
Representative plaque assay images. Representative images of a single well of a 6-well plate shown for each candidate grown in ±IAA.

**Figure S6:**
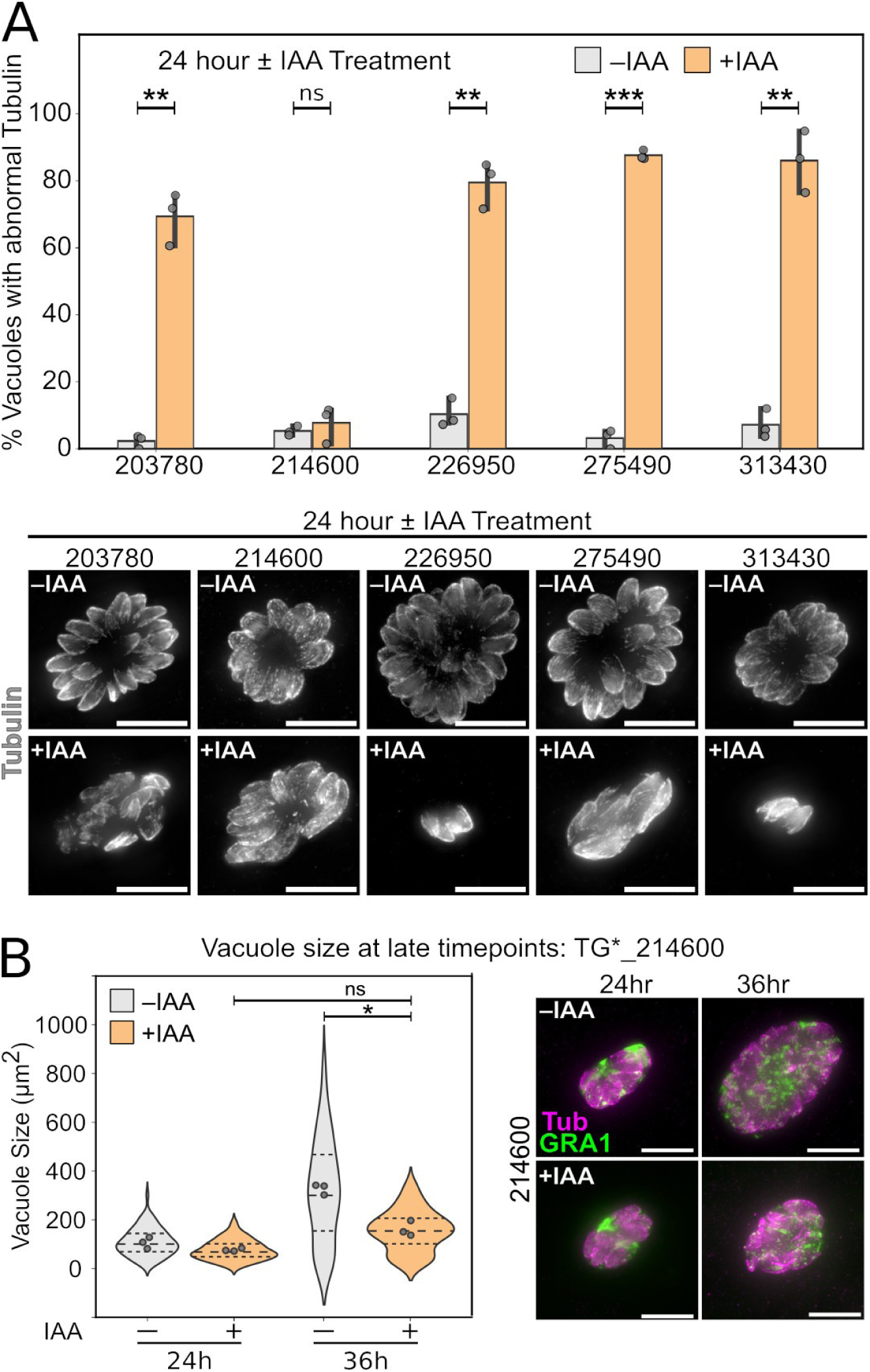
Block in replication at late timepoints for a subset of essential Nups. (A) The percent of vacuoles showing abnormal tubulin staining after 24 h growth in ±IAA. Note that while TG*_214600 shows no quantifiable phenotype, tubulin staining does not appear entirely normal after 24 h growth in +IAA. (B) TG*_214600 AID parasites grown in +IAA showed no significant increase in vacuole size from 24-36h, suggesting a block in growth at late timepoints. All scale bars are 10 μm.

**Figure S7:**
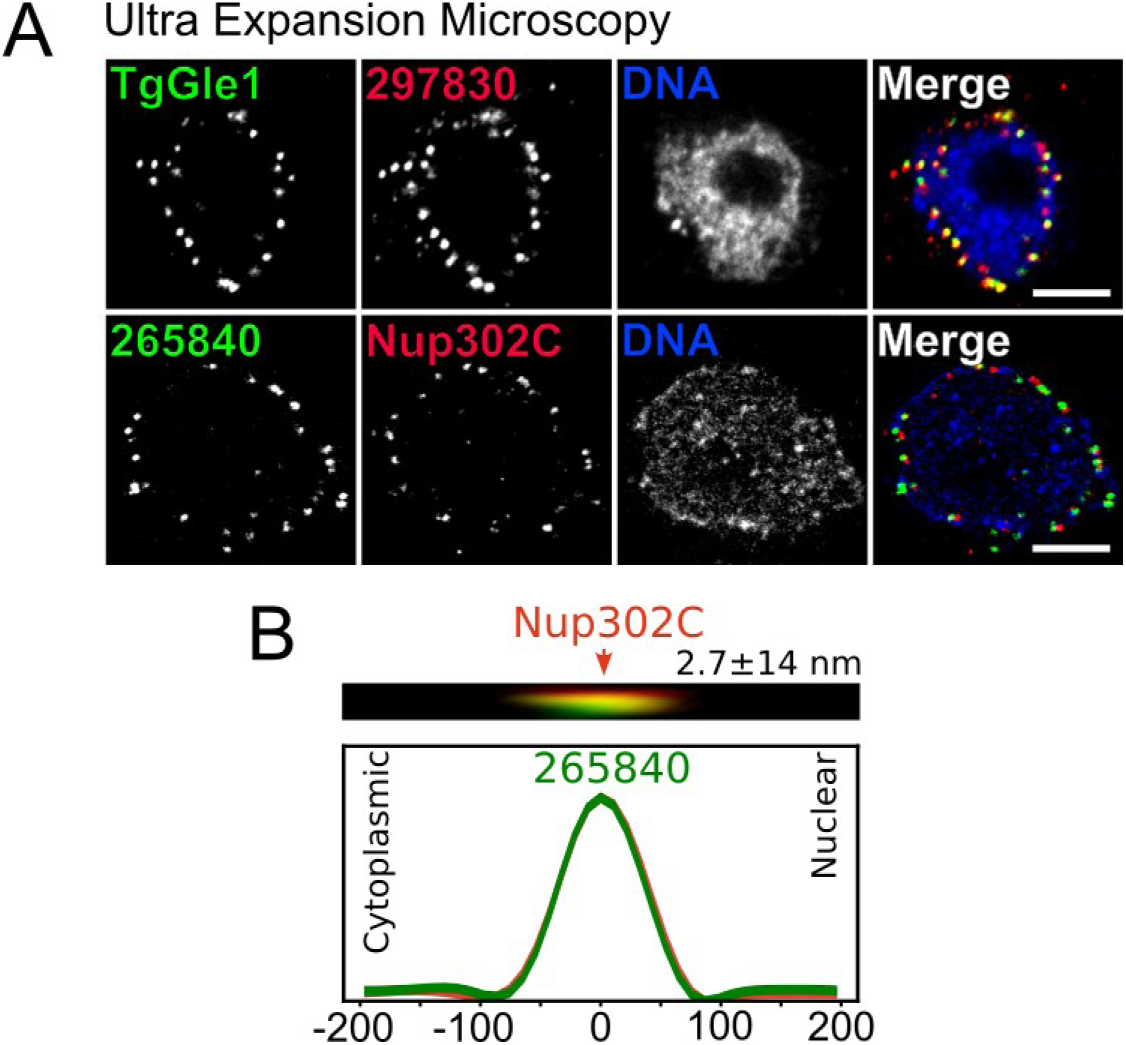
UExM of Gle1/TG*_259172 and Nup148/TG*_265840. (A) Maximum intensity projections of 10 slices of a UExM confocal stack taken from the middle of the parasite nucleus. All scale bars are 5 μm. (B) Representative line scan profile used to measure distance between TG*_265840 puncta and NPC marker Nup302C. A representative image for the line scan is above the plot. Distance is mean ± sem from n=10 puncta.

**Figure S8:**
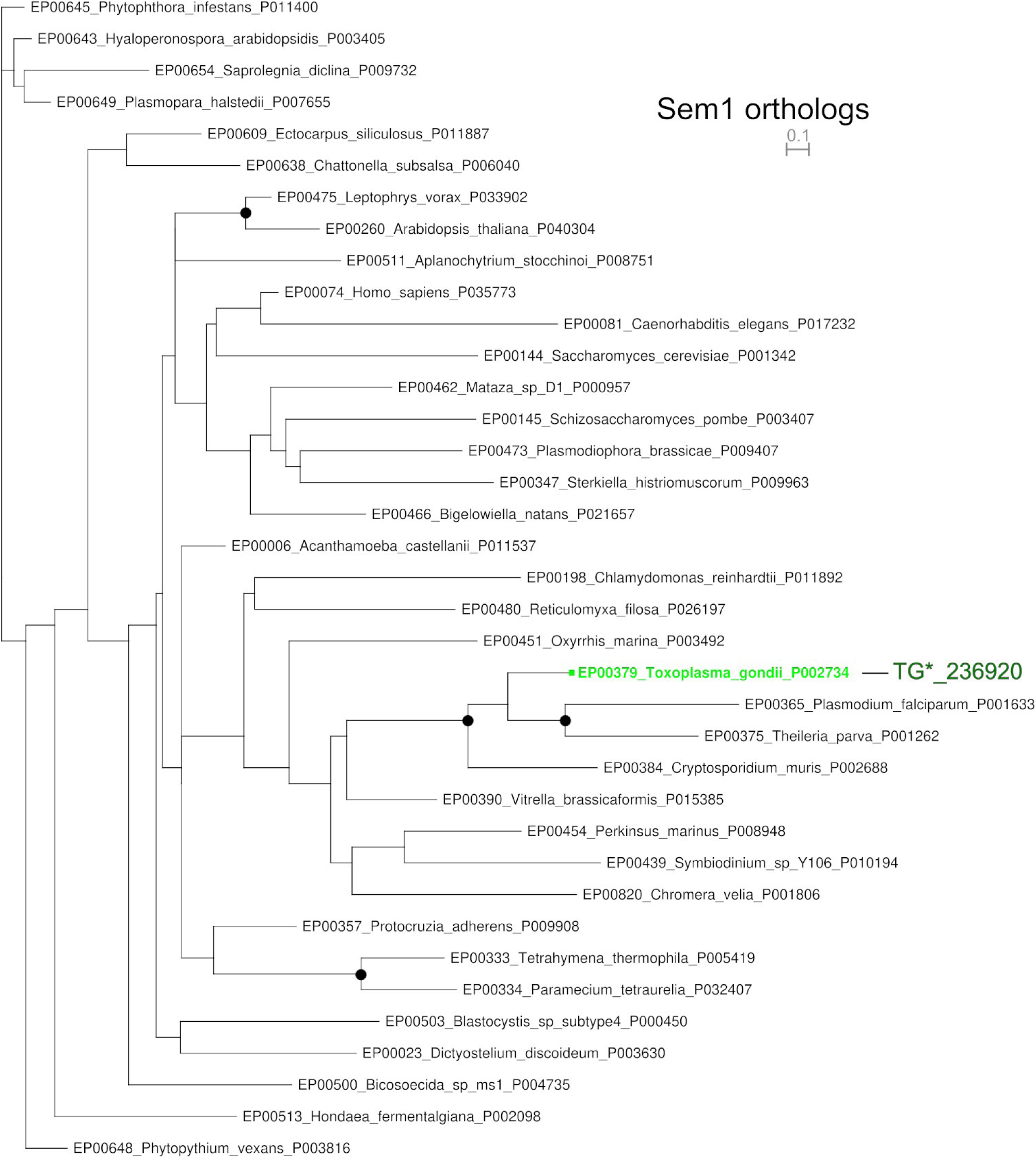
Phylogenetic tree of Sem1 orthologs. Phylogenetic tree of Sem1 orthologs from diverse eukaryotes show potential members in *Toxoplasma* (green) and other members of the Alveolate and SAR clade. Black circles indicate >95% bootstrap support. Tree with all support values and branch lengths is available in Data S4. Accession numbers are in Data S5. Scale bar is substitutions per position.

**Figure S9:**
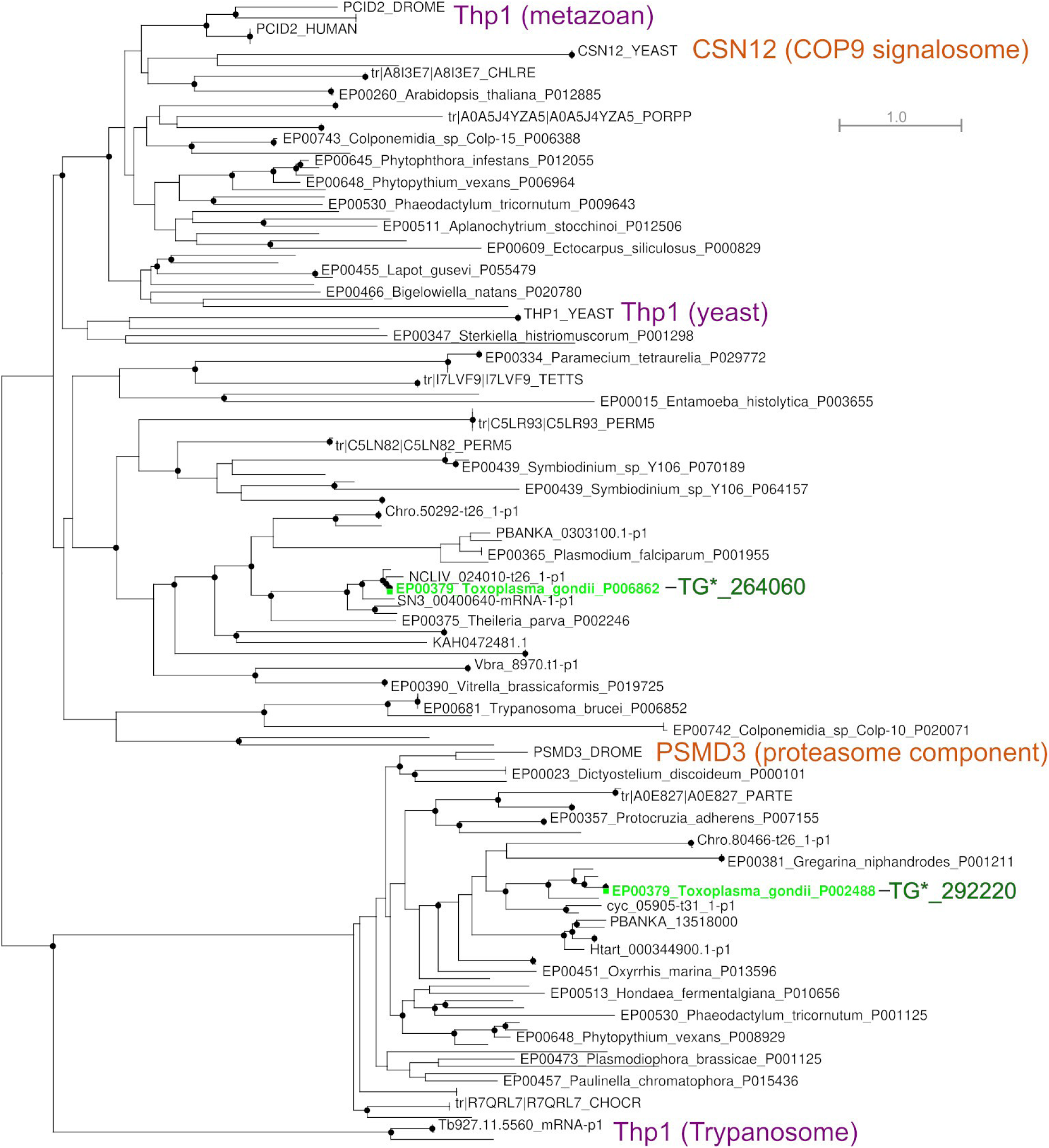
Phylogenetic tree of Thp1 orthologs. Phylogenetic tree of potential orthologs of Thp1 and its paralog PSMD3 (a proteasomal subunit) and CSN12 (a COP9 signalosome component) from diverse eukaryotes show potential members in *Toxoplasma* (green) and other members of the Alveolate and SAR clade. Black circles indicate >95% bootstrap support. Tree with all support values and branch lengths is available in Data S4. Accession

**Figure S10:**
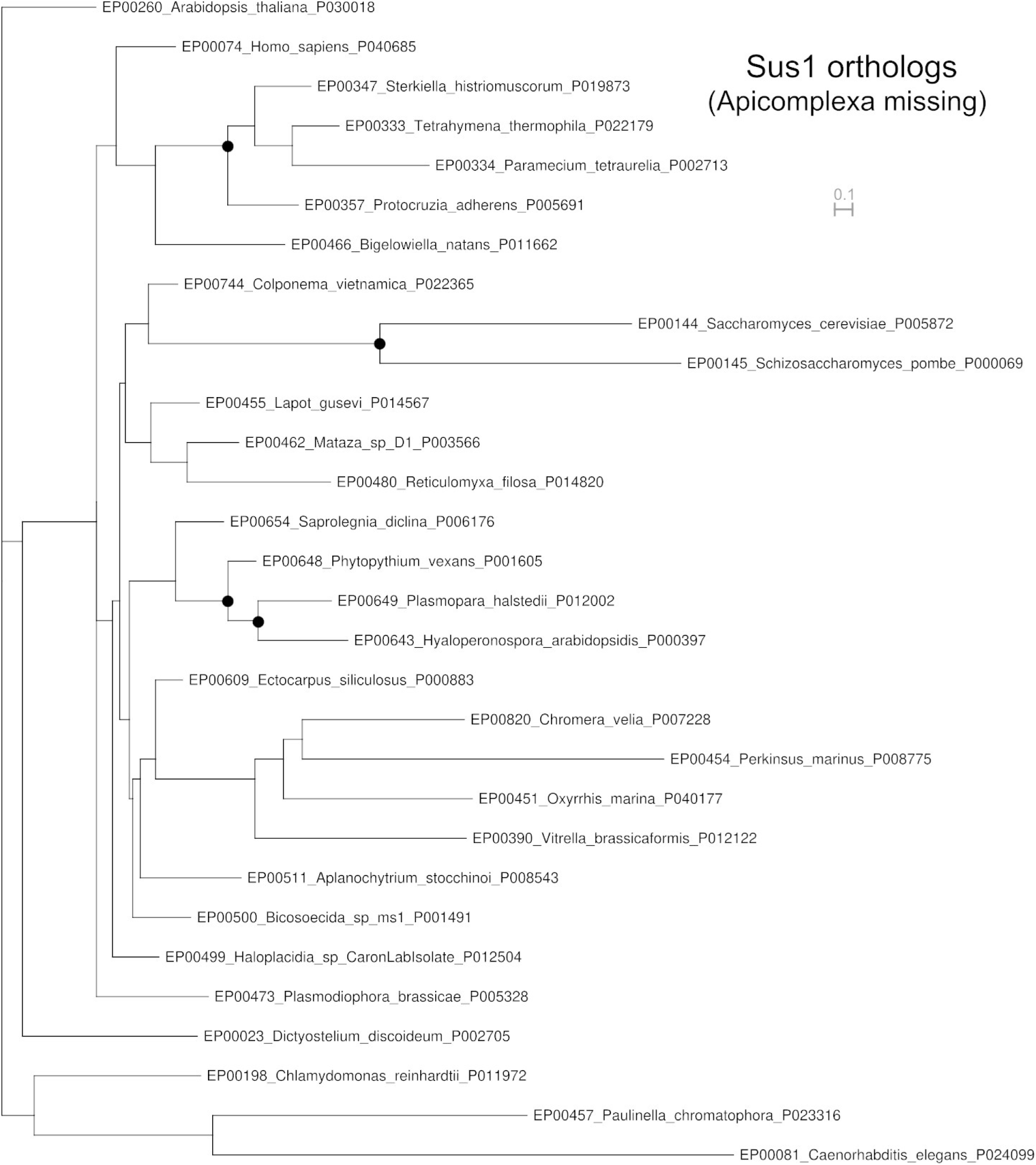
Phylogenetic tree of Sus1 orthologs. Phylogenetic tree of potential orthologs of Sus1 from diverse eukaryotes show throughout Alveolates and other members of the SAR clade but no candidate orthologs from Apicomplexa. Black circles indicate >95% bootstrap support. Tree with all support values and branch lengths is available in Data S4. Accession

**Figure S11:**
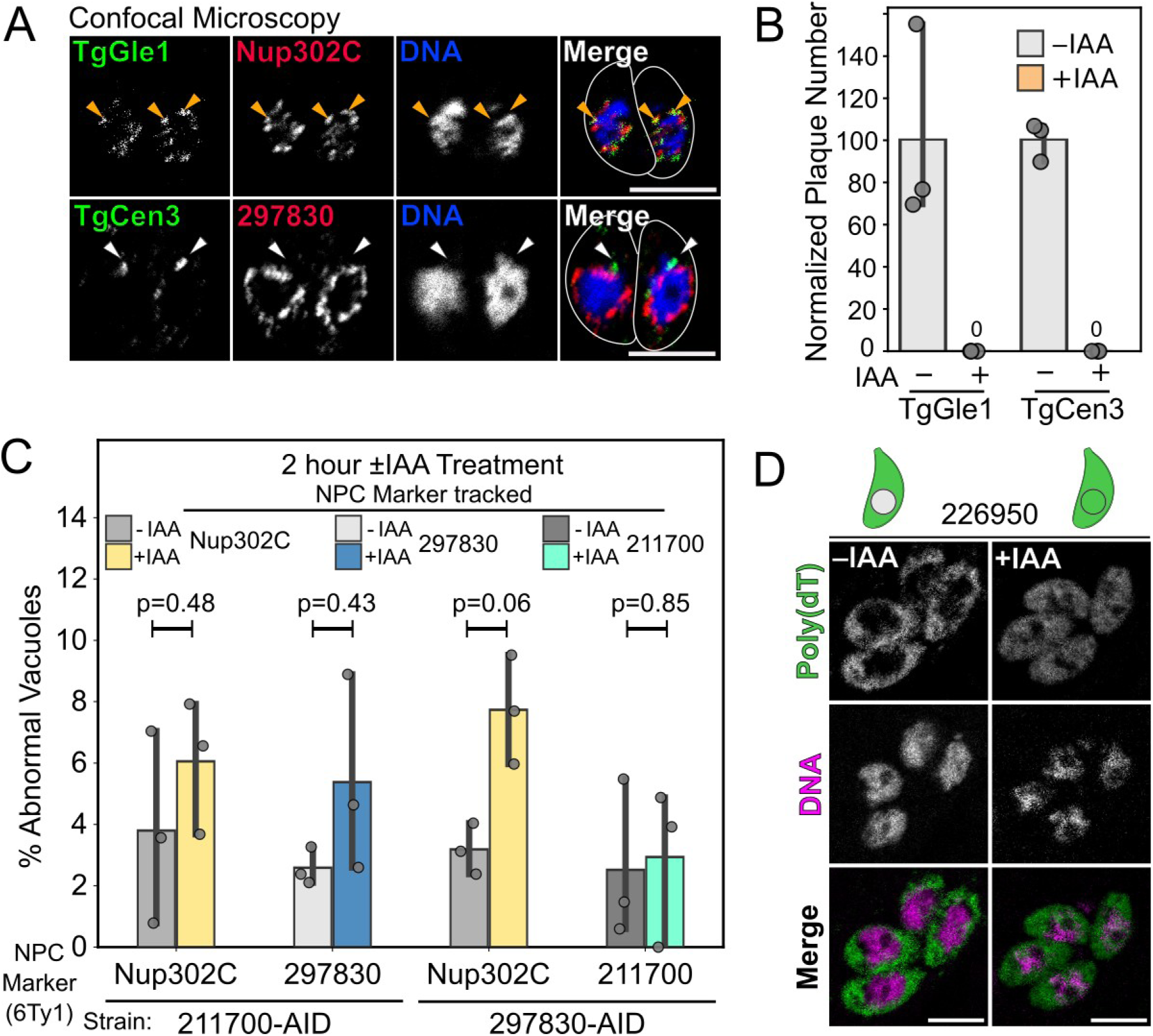
Characterization of additional proteins involved in mRNA export. (A) TgGle1^AID^ localizes to the NPC (orange arrows), while TgCen3AID-3xHA shows expected punctate staining consistent with centrosomal localization (white arrows). Scale 5µm. (B) Both TgGle1^AID^ and TgCen3^AID^ strains are unable to form plaques when grown in +IAA. (C) Quantification of the indicated markers (Nup302C, 297830, or 211700) in TG*_211700^AID^ and TG*_297830^AID^ strains after incubation in ±IAA for 2 hours. p-values (non-significant) indicated from unpaired two-tailed Student’s t-test. (D) Confocal slices of TG*_226950^AID^ parasites that were allowed to infect overnight and then grown in +IAA media for 2 hours before fixation. mRNA was stained by FISH with poly(dT) oligo (green) and parasites were counterstained with Hoechst (magenta). Cartoon parasites represents healthy parasites next to -IAA and parasites with mRNA export block next to +IAA.

**Figure S12:**
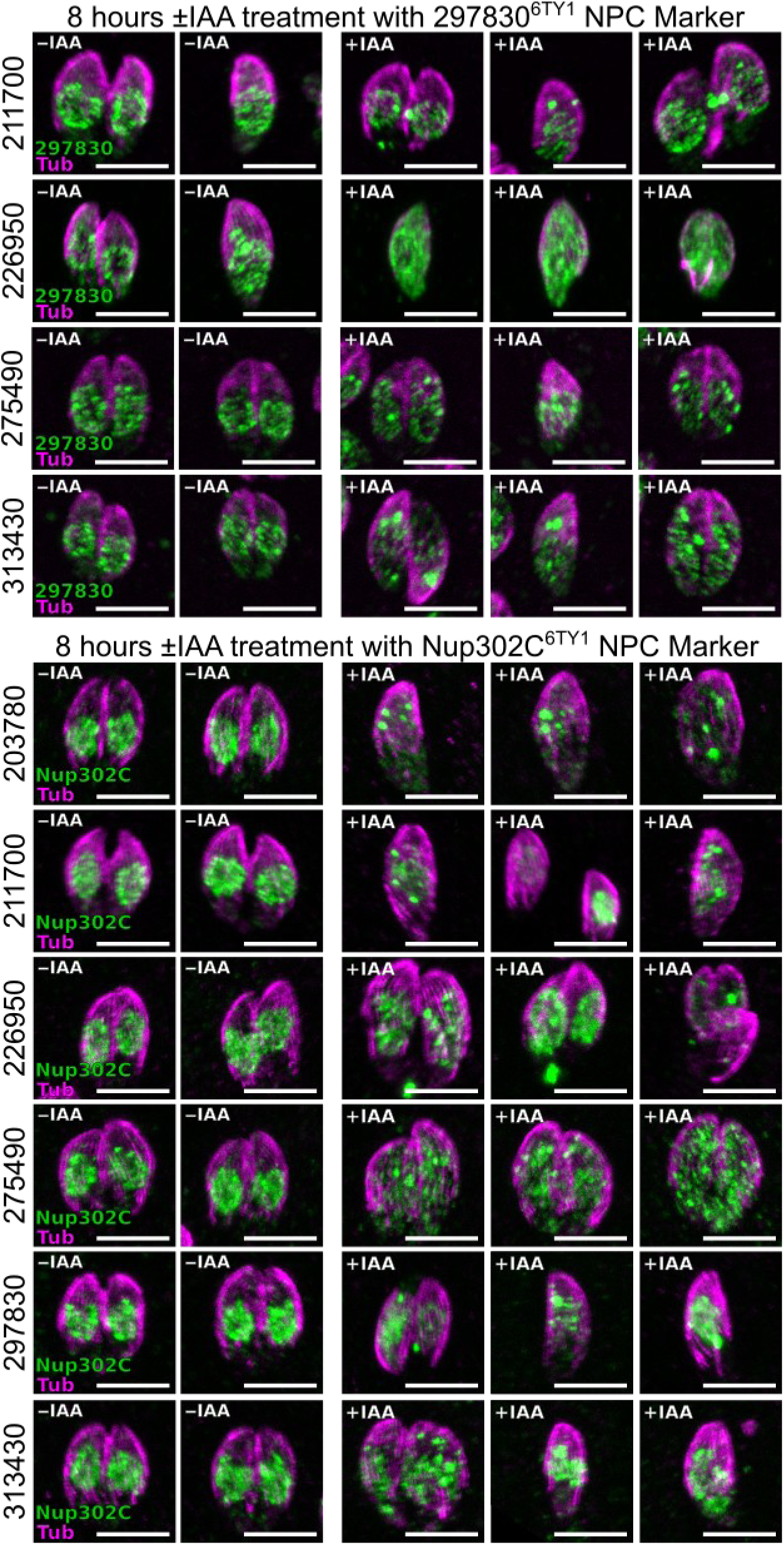
Mislocalization of NPC markers seen in parasites upon knockdown of Nups. Maximum intensity projections of confocal stacks of parasites grown 8 h in ±IAA media. These images are representative of the images used to quantify mislocalization of NPC markers in Figure 8C. Scale bars are 5 µm.

**Figure S13:**
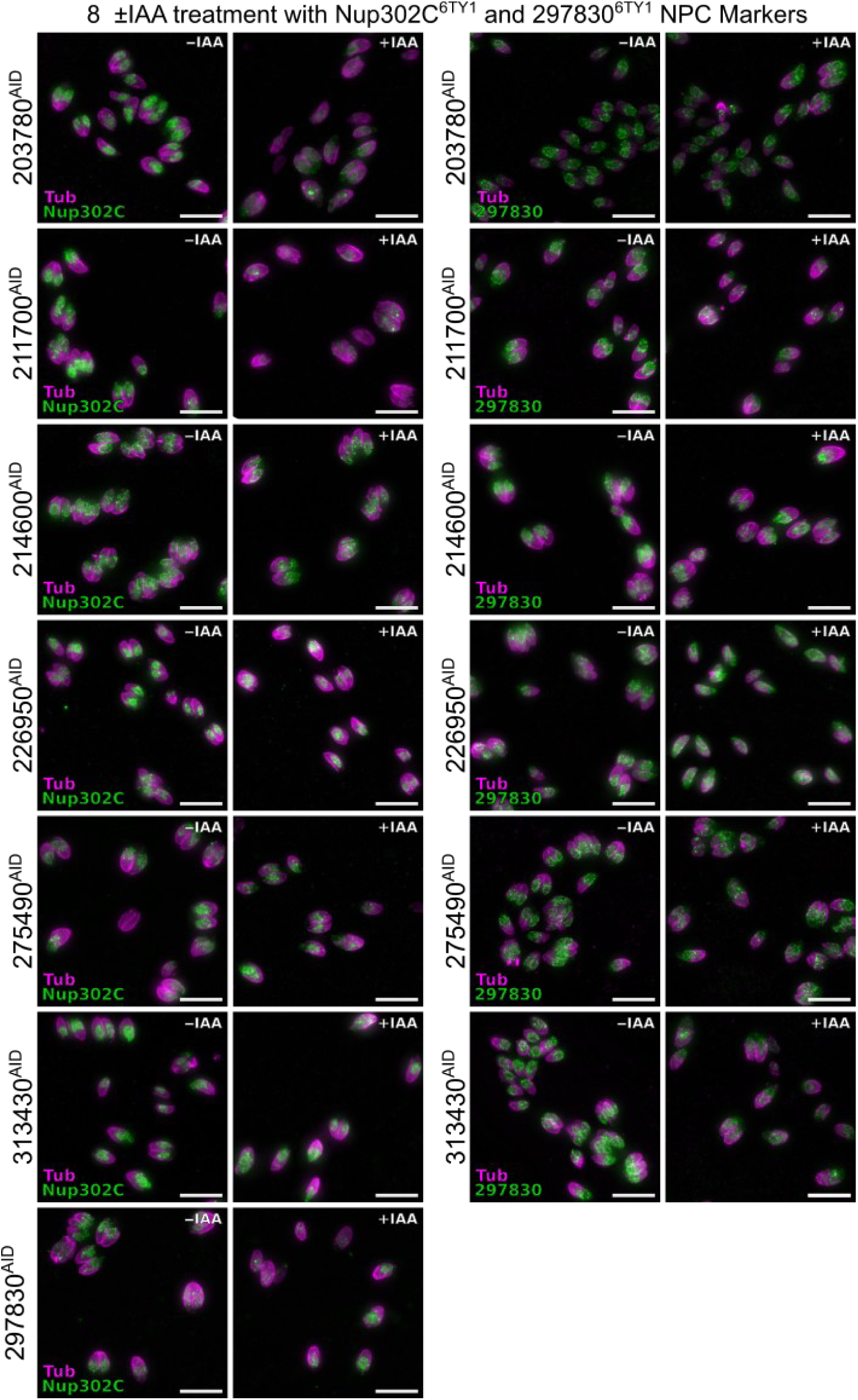
Representative images used in the quantification of NPC marker mislocalizaiton. Maximum intensity projections of widefield stacks of parasites grown 8 h in ±IAA media. These images are representative of the images used to quantify mislocalization of NPC markers in Figure 8C. Scale bars are 10 µm.

**Figure S14:**
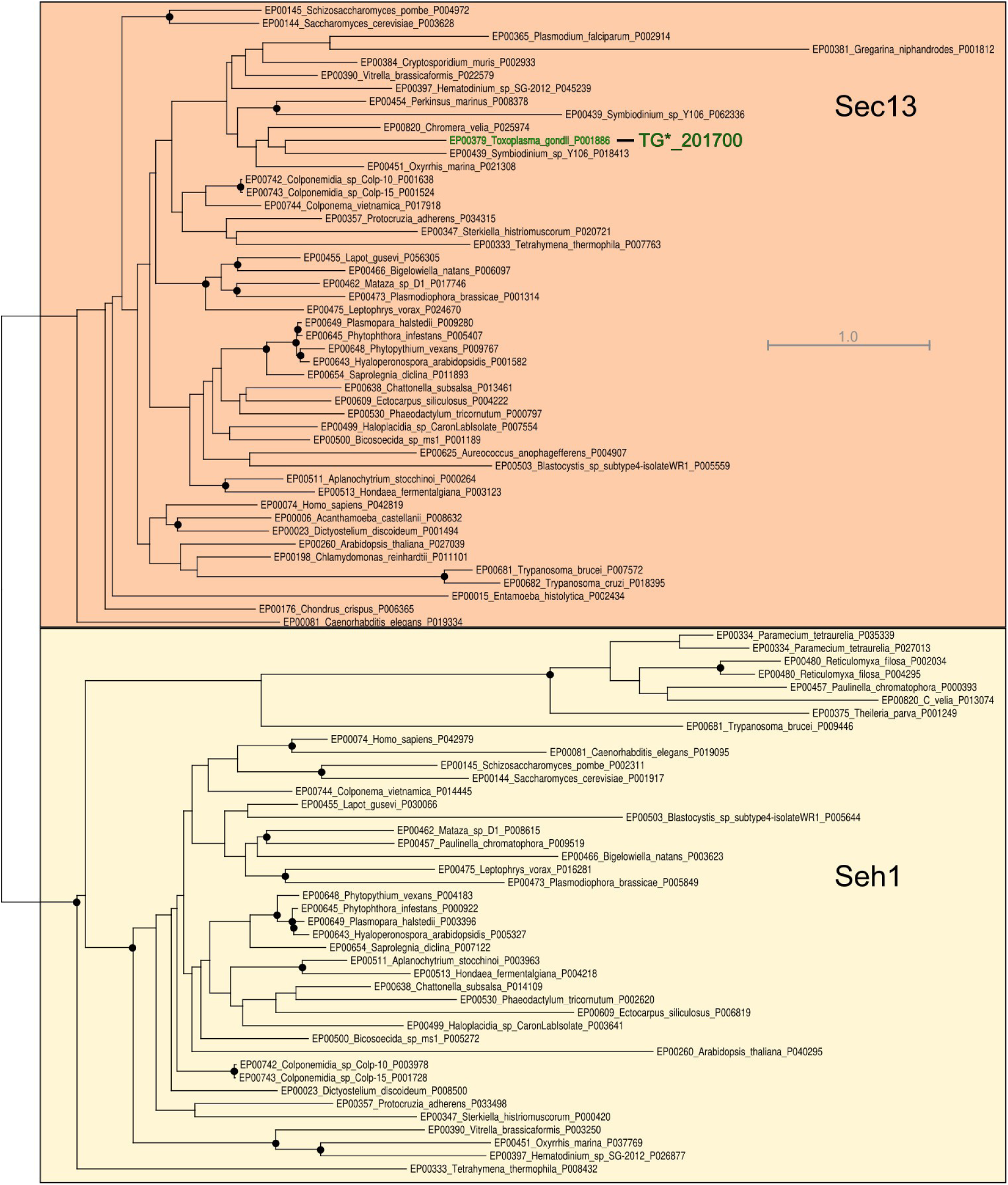
Phylogenetic tree of Sec13/Seh1 orthologs. Phylogenetic tree of potential orthologs of Sec13 and its Nup paralog Seh1 from diverse eukaryotes shows Sec13 is conserved in diverse Alveolates. Seh1, however, is found in ciliates appears to be missing in Apicomplexa and dinoflagellates. Black circles indicate >95% bootstrap support. Tree with all support values and branch lengths is available in Data S4. Accession numbers are in Data S5. Scale bar is substitutions per position.

**Figure S14:**
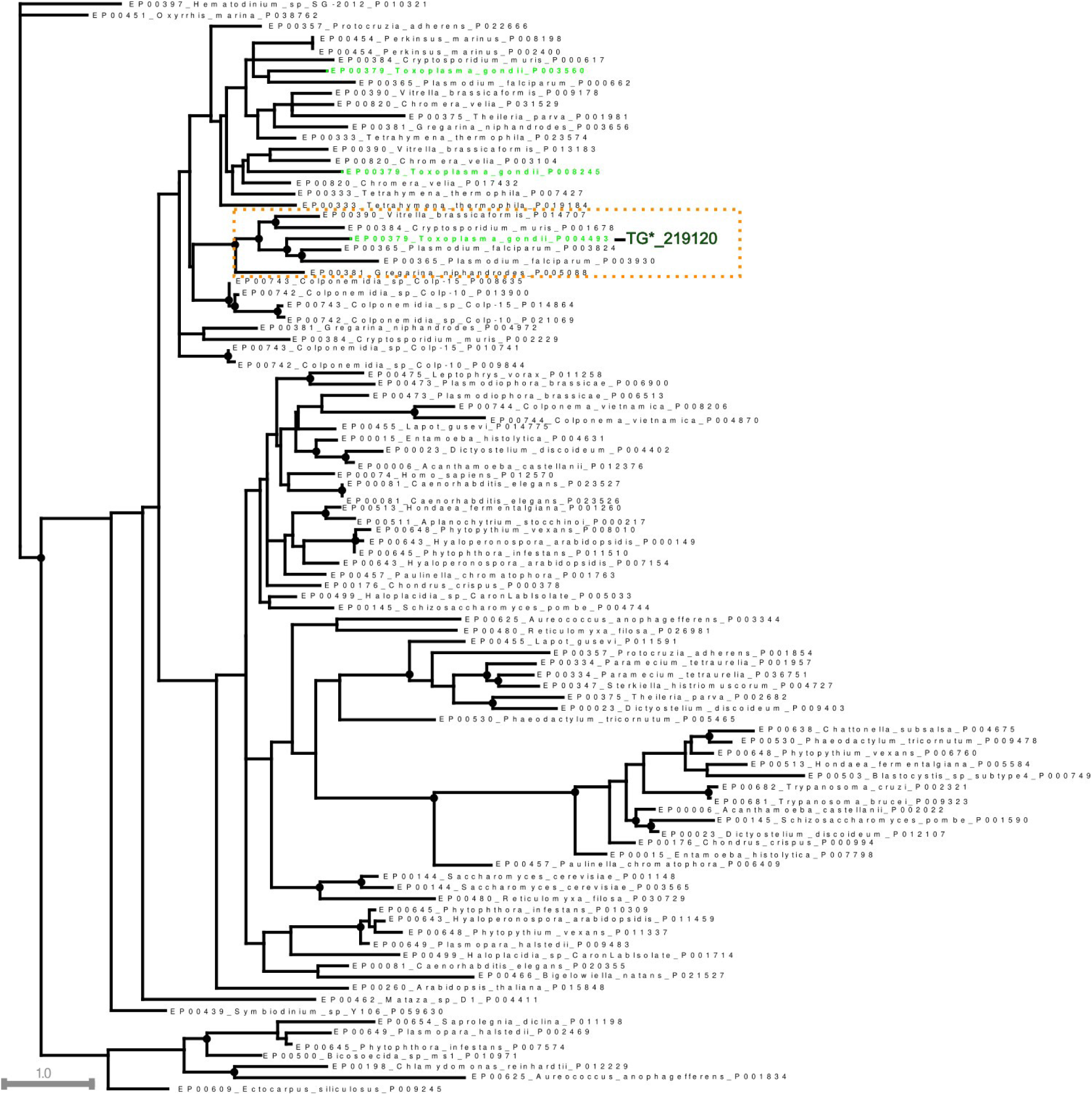
Phylogenetic tree of Zn-finger proteins related to TG*_219120. Phylogenetic tree of Zn-finger domains with strong homology to TG*_219120. While TG*_219120 is conserved throughout Apicomplexa, this subfamily seems limited to these parasites. *Toxoplasma* has two other paralogs (green) that are likely nuclear regulators of mRNA decay. Black circles indicate >95% bootstrap support. Tree with all support values and branch lengths is available in Data S4. Accession numbers are in Data S5. Scale bar is substitutions per position.

## Supplemental Data Details

**Supplemental Data S1:** Mass spectrometry data from proximity biotinylation experiments. Mass spectrometry data are available from the MassIVE database with accession MSV000098791.

**Supplemental Data S2:** Spreadsheet detailing search hits and corresponding e-values from OrthoMCL, BLAST, PANTHER, and Foldseek.

**Supplemental Data S3:** List of accession numbers used in phylogenetic analyses.

**Supplemental Data S4:** Complete alignments and phylogenetic trees used in this study.

**Supplemental Data S5:** List of primer sequences used in this study.

